# Two decades of suspect evidence for adaptive molecular evolution – Negative selection confounding positive selection signals

**DOI:** 10.1101/2021.11.22.469483

**Authors:** Qipian Chen, Hao Yang, Xiao Feng, Qingjian Chen, Suhua Shi, Chung-I Wu, Ziwen He

## Abstract

There is a large literature in the last two decades affirming adaptive DNA sequences evolution between species. The main lines of evidence are from i) the McDonald-Kreitman (MK) test, which compares divergence and polymorphism data, and ii) the PAML test, which analyzes multi-species divergence data. Here, we apply these two tests concurrently on the genomic data of *Drosophila* and *Arabidopsis*. To our surprise, the >100 genes identified by the two tests do not overlap beyond random expectation. Because the non-concordance could be due to low powers leading to high false-negatives, we merge every 20 - 30 genes into a “supergene”. At the supergene level, the power of detection is large but the calls still do not overlap. We rule out methodological reasons for the non-concordance. In particular, extensive simulations fail to find scenarios whereby positive selection can only be detected by either MK or PAML, but not both. Since molecular evolution is governed by positive and negative selection concurrently, a fundamental assumption for estimating one (say, positive selection) is that the other is constant. However, in a broad survey of primates, birds, *Drosophila* and *Arabidopsis*, we found that negative selection rarely stays constant for long in evolution. As a consequence, the variation in negative selection is often mis-construed as signals of positive selection. In conclusion, MK, PAML or any method that examines genomic sequence evolution has to explicitly address the variation in negative selection before estimating positive selection. In a companion study, we propose a possible path forward in two stages – first, by mapping out the changes in negative selection and then using this map to estimate positive selection. For now, the large literature on positive selection between species has to await the re-assessment.

## Introduction

The inferences of adaptive evolution in DNA sequences permit the assessment of the biological significance of genes of interest. Such inferences may then guide the planning of functional validation. Extensive reports of adaptively evolving genes can be found in almost all taxa [1–3] as well as all types of cancers [4,5]. Indeed, the large-scale genomic data amassed in the last two decades have led to the acceptance of pervasive adaptive evolution over the neutral theory of molecular evolution [3,6–9].

The detection of positive selection largely falls into two broad classes [10–14]. One class attempts to detect positive selection that operates within populations [10,13,15]. The other focuses on positive selection that operates in the longer term, i.e., the divergence between species [16–18]. Methods of either class may use data of both polymorphism and divergence [12,16,19]. Positive selection signals could be abundant between species but undetectable within populations, or vice versa. It is hence possible to reject the neutral theory in part (either within or between species) or in whole.

In this study, we focus on positive selection between species. The results of analyses between species can be qualitatively different from the analyses of polymorphism data within species. The two approaches are complementary, rather than redundant (see Discussion) [20–22]. There are several challenges in correctly inferring positive selection since DNA sequences are simultaneous influenced by multiple forces that may include mutation, genetic drift, positive selection and negative selection. To tease apart these forces often requires making assumptions about “other” forces. In particular, it is usually assumed that negative selection is constant in the time frame of interest, often in tens of millions of years.

In between-species tests, one compares the number of non-synonymous changes per non-synonymous site (Ka or d_N_) with the per-site synonymous changes (Ks or d_S_) [11,17,23]. The Ka/Ks (or d_N_/d_S_) ratio will deviate from 1 if non-synonymous changes are under stronger selection than synonymous substitutions. In the absence of selection, *R* = Ka/Ks ∼ 1, which is the hallmark of neutral evolution [24,25]. In among-species comparisons, genome-wide *R* ranges mainly between 0.05 and 0.25 [25,26], thus indicating the prevalence of negative selection. When *R* > 1, positive selection is evident. However, *R* > 1 is too stringent a criterion as it requires positive selection to overwhelm negative selection. Indeed, few genes in any genome comparison have *R* significantly greater than 1 [14,27].

The two commonly used methods that relax the requirement for *R* > 1 over the entire gene are the MK (McDonald-Kreitman) [12,16] and the PAML (Phylogenetic analysis by maximum likelihood) [28,29] tests. More recent tests, such as those given in Ref. [21,30–32], are not included because they have been used far less frequently. If the detected adaptive signals are true, the results from the two tests are expected to show substantial overlap. However, since their proposals and after extensive applications, the two tests have rarely been used side-by-side. If the results from the two types of analyses are non-concordant, half or even much of the large literature on positive selection at the genomic level may have to be cast in doubt.

### Theoretical bases of the MK and PAML tests

While Ka and Ks are the cornerstones for detecting natural selection in coding sequences, they can only inform about either positive or negative selection, but not both. This is because Ka/Ks, when averaged over all sites, is the joint outcome of the two opposing forces, as described by the basic population genetic theory:

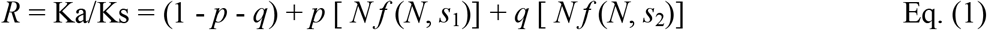

For simplicity, we consider an ideal population of haploids with size *N*. (For a diploid population, *N* should be replaced by 2*N*.) In Eq. (1), *p* and *q* are the proportion of advantageous and deleterious mutations, respectively [24,33,34]. Also, *f* (*N*, s) = (1 - *e*^-2*s*^)/ (1 - *e*^-2*Ns*^) is the fixation probability of a mutation with a selective coefficient *s* that can be > 0 (denoted by *s*_1_) or < 0 (*s*_2_) [34,35]. Both *s*_1_ and |*s*_2_| are assumed to be larger than 1/2*N*, below which selection is too weak to matter. If *s*_1_ is small (but no smaller than 1/2*N*), then *f* (*N, s*_1_) ∼ 2*s*_1_. Similarly, if |*s*_2_| >> 1/2*N, f* (*N, s*_2_) ∼ 0, meaning no fixation of deleterious mutations. Eq. (1) is then reduced to

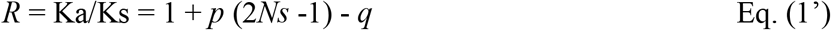

Following Eq. (1’), if Ka/Ks = 0.2, for example, the null hypothesis of neutrality would assume *p* = 0 and *q* = 0.8 with 20% neutral mutations. However, it is also possible that *p* and *q* could both be larger, for example, *p* = 0.01, 2*Ns* = 11 and *q* = 0.9, yielding the same Ka/Ks = 0.2.

A central task of molecular evolutionary studies is to estimate *p* (and, when possible, *Ns*) from a set of DNA sequences within and between species. Two typical examples that are used in this study can be found in Fig. 1a and 1b, from species in the *Drosophila* and *Arabidopsis* clade, respectively (Fig. 1). In order to estimate *p* using Eq. (1’), one would have to know the value of *q*. However, the question is whether *q* can in fact be estimated, for example, if *q* fluctuates in time. The field deals with this question not by answering it but by assuming that *q* is constant in time and across lineages (and can indeed be estimated). The difference between MK and PAML, as discussed below, is hence in how to estimate *q*. Broadly speaking, MK estimates *q* explicitly from the polymorphism of a single species (the blue triangles in Fig. 1a and 1b) but PAML takes the average of *q* across the entire phylogeny. If *q* is constant, both approaches are valid. Comparing the results of MK and PAML would amount to testing the common assumption that *q* is constant.

**Figure 1.**
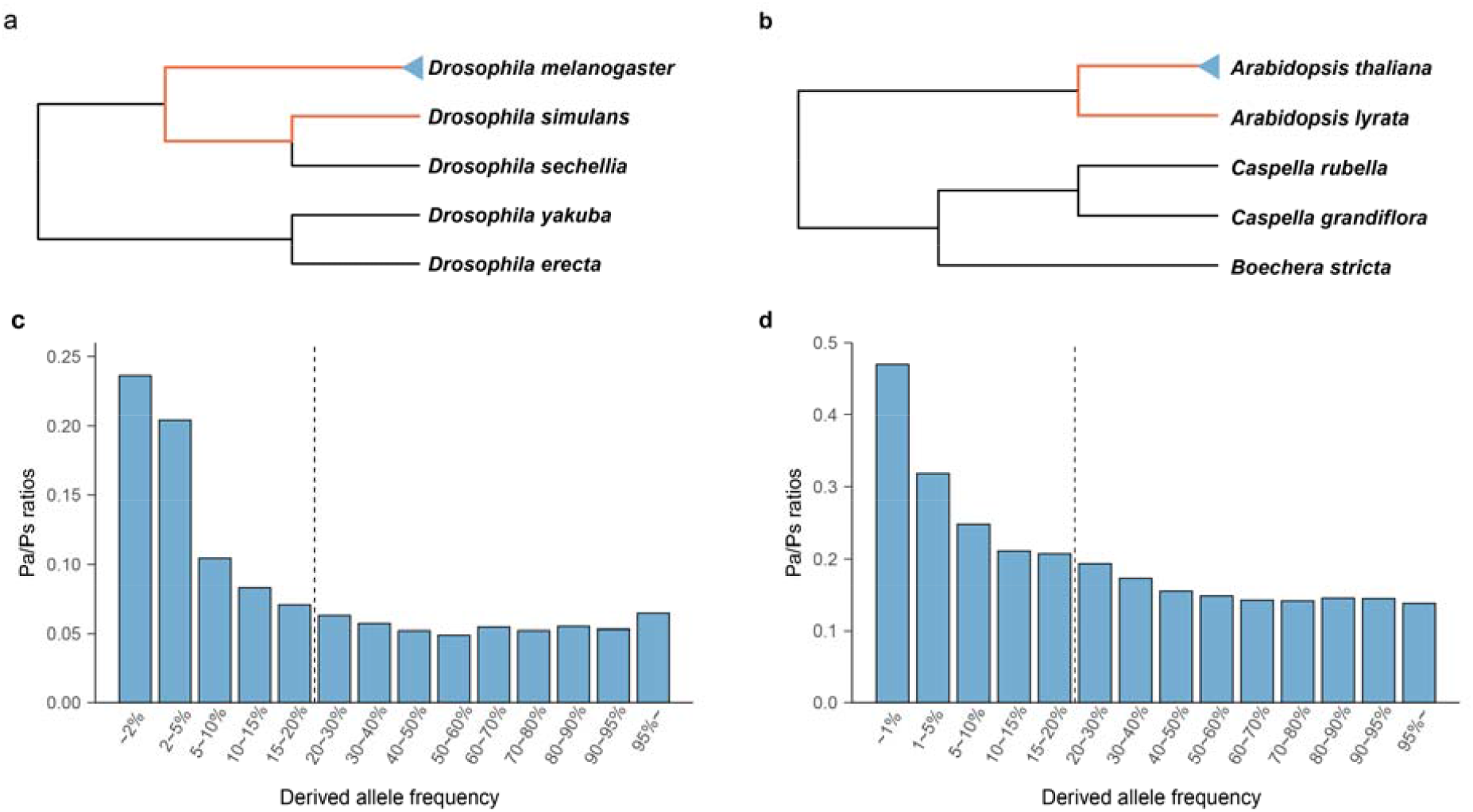
Between-species divergence and within-species polymorphism for detecting positive selection. **a** and **b**, Phylogeny of *Drosophila* and *Arabidopsis* species. Both the MK and PAML tests are forced to detect positive selection along the branches marked by red. The MK test uses polymorphisms (indicated by the blue triangle) for reference. The reference for PAML is described in Supplementary Methods. **c** and **d**, The Pa/Ps ratio as a function of the mutant frequency in *D. melanogaster* and *A. thaliana*. The dashed line, separating low- and high-frequency bins, is placed where the Pa/Ps ratio reaches a steady level.

The MK test is usually applied to a particular phylogenetic lineage, marked by the red line in Fig. 1a and 1b. The Ka and Ks values in the red line lineage are contrasted with the corresponding polymorphisms (Pa and Ps) in the blue triangle. The value of Pa and Ps denotes, respectively, the level of non-synonymous and synonymous polymorphism (per site) within a species. The rationale of the MK test is that *p* ∼ 0 in the polymorphism data thanks to the rapidity with which advantageous mutations are fixed. Thus, Eq. (1) becomes

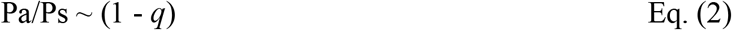

The *q* [ *N f* (*N, s*_2_)] term of Eq. (1) should be treated differently in Eq. (2) because deleterious mutations are common in the polymorphism data, but mainly in the low-frequency range. To use Eq. (2), rigorous data processing is necessary to remove deleterious mutations (see Fig. 1c and 1d). In short, the MK test estimates *q* from Eq. (2) and applies it to extracts *p* from Eq. (1’).

In contrast to MK, PAML does not explicitly estimate *q*. PAML compares the substitution numbers across many lineages to identify positively (or negatively) selected genes on the assumption that unusually high (or low) numbers could be indicative of selection. In particular, the proportion of adaptive sites that have a higher non-synonymous than neutral rate is estimated. Neither does PAML use polymorphic data to supplement the estimation of *q*. There are three (sub-) models in PAML, each representing a different set of assumptions. The site model identifies sites with an increase or decrease in non-synonymous substitutions in the entire phylogeny [18,28,29]. The branch-site model compares sites of a pre-selected branch (the foreground) to other sites on all branches as well as the same sites on other branches (the background) [8,36,37]. The third sub-model is not considered here.

## Results

Despite the very different approaches, the MK and PAML tests can be used to answer the same question – How much adaptive evolution has happened in the chosen genes on a given branch (e.g., the red-line branch of Fig. 1a and 1b)? As stated above, the concordance between the two tests is the best way to check the validity of the extensive literature on adaptive DNA evolution.

Because the MK test is about positive selection along the red line, it does not offer any information about selection elsewhere in the phylogeny. Therefore, it is necessary to compare it to each of the two PAML sub-models. If the MK test identifies genes that are generally prone to adaptive evolution, the proper comparison would be the PAML site model. Alternatively, if the adaptation is specific to a specific branch, then the branch-site model would be a more suitable comparison. We will present the site model results in the main text and the branch-site model results in Supplementary Information. The two sets of comparisons lead to the same conclusion, although the site model appears to be statistically more robust.

PART I tests the concordance between MK and PAML on *Drosophila* and *Arabidopsis* data. PART II examines (and rejects) the methodological explanations. PART III provides evidence for a biological explanation based on non-constant negative selection.

### PART I. Comparing MK and PAML test results in *Drosophila* and *Arabidopsis*

#### 1. Identifying adaptive genes with high stringency

For a quick overview of the absolute and relative performances of MK and PAML, we first present the distribution of the P value across genes. The MK test P values were obtained from Fisher’s exact test site count contingency tables. The likelihood ratio test was used to obtain PAML P values. The P value distributions are shown in the four panels of Fig. 2 for two taxa and two tests. The distribution is concentrated above P = 0.8 (the MK test for *Drosophila*) and P = 0.9 (the other panels). This concentration means that a very large percentage of genes show no detectable signal, partly because most genes experience too few changes to be statistically informative. Furthermore, the null model does not fully incorporate factors that can affect the test. For example, the polymorphism data may not reflect the complete removal of deleterious mutations, and the strength of negative selection is often under-estimated [3,38,39].

**Figure 2.**
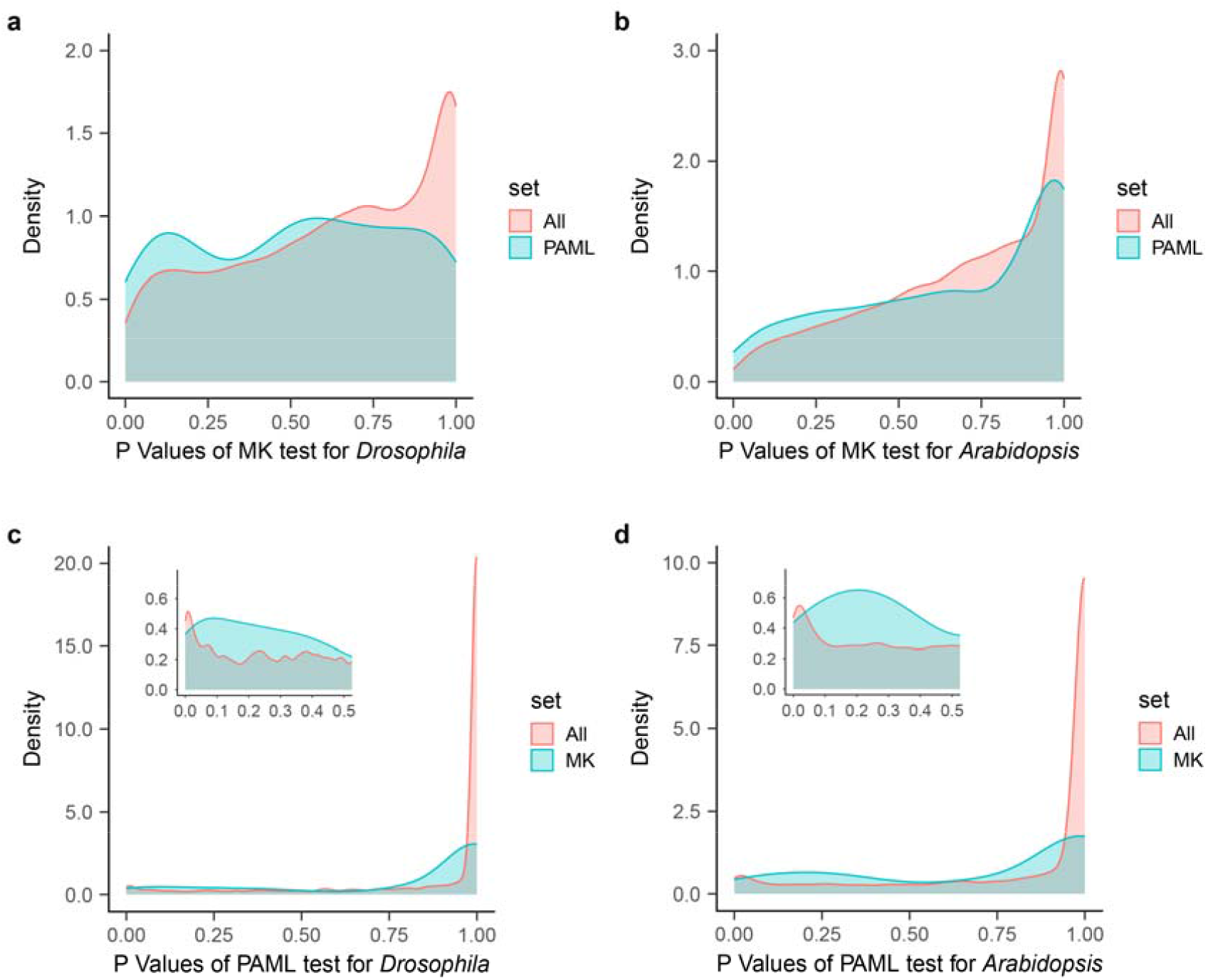
P value distributions of the MK and PAML test. **a** and **b**, P values of the MK test for *Drosophila* and *Arabidopsis*. The distribution for all genes is shown in red, and the distribution for genes pre-filtered by the PAML test is shown in blue. **c** and **d**, P values of the PAML test. Results of genes pre-filtered by the MK test are shown in blue. These two panels are the mirror images of panels (a-b) with MK and PAML switched.

Fig. 2 also shows that far fewer than 5% of them would be detected as adaptive at the 5% cutoff. We therefore compare the observed P values from the MK and PAML tests against each other rather than against the null model. In each panel of Fig. 2, one line represents the test results on all genes, and the other is derived from loci that have been pre-filtered through the other test. In Fig. 2a-2b, genes pre-filtered through PAML have smaller P values in the MK test, reflected by the leftward shift in the P value distribution. The same is true in Fig. 2c-2d, where pre-filtering by MK reduces the PAML test P values. The two tests are indeed correlated, but only weakly. This is also true in Fig. S1, where the branch-site model of PAML is used.

Knowing the weak correlation between the two tests, we enumerate the overlap between them by comparing the candidate adaptive genes with P < 0.05. Given the P value distributions shown in Fig. 2, these genes are merely the most likely candidates proposed by each test. Hence, significant overlaps would be mutual corroborations. For the “individual genes” analysis in *Drosophila*, we identified 186 from 5425 genes by the MK test and 145 genes by PAML, corresponding to 3.43% and 2.67% of the genome (see Table 1). The overlap between these two sets contains only nine genes. Although the observed overlap is higher than the expected 4.97 (P < 0.1, Fisher’s exact test), the overlap is too small to be biologically meaningful. The same pattern is true for *Arabidopsis*, in which 145 and 505 genes are called by these two tests, but only 14 genes are called by both tests. Again, the observed overlap is significantly higher than the expected 5.55 (P < 0.01, Fisher’s exact test), but the actual overlap is minimal. A simple explanation for the non-overlap is a high false-negative rate. In other words, each test may have detected only a small fraction of the true adaptive genes.

**Table 1.** Proportion of adaptively evolving genes identified by two tests (P < 0.05)

| **Gene Category** | **MK** | **PAML** | **Expected overlap** | **Observed overlap** |
| --- | --- | --- | --- | --- |
| *Drosophila* |  |  |  |  |
| Individual genes | **3.43%**<br>(186/5425) | **2.67%**<br>(145/5425) | **0.09%**<br>(4.97/5425) | **0.17%**<br>(9/5425) |
| Supergenes<sup>a</sup> | **56.0%**<br>(112/200) | **18.0%**<br>(36/200) | **10.1%** | **10.0%**<br>(20/200) |
| Component genes<sup>b</sup> | **5.04%**<br>(158/3132) | **5.77%**<br>(60/1040) | **0.29%** | **0.48%**<br>(3/619) |
| *Arabidopsis* |  |  |  |  |
| Individual genes | **1.12%**<br>(145/12975) | **3.89%**<br>(505/12975) | **0.04%**<br>(5.55/12975) | **0.11%**<br>(14/12975) |
| Supergenes | **8.20%**<br>(41/500) | **25.6%**<br>(128/500) | **2.10%** | **2.00%**<br>(10/500) |
| Component genes | **3.63%**<br>(38/1048) | **7.44%**<br>(246/3306) | **0.27%** | **0.78%**<br>(2/258) |

<sup>a</sup> Supergenes are concatenations of 20-30 neighboring genes by physical location. See Table S1 for supergenes concatenated by gene function.
<sup>b</sup> Component genes are individual genes within supergenes that have passed the MK and/or PAML test.

#### 2. The analysis of supergenes and their component genes

Since a gene on average harbors only a few substitutions, the power to reject the null model is often low. To augment the statistical power, we created artificial “supergenes” by merging 20 to 30 genes into a longer sequence. In the statistical sense, a supergene is like any individual gene that comprises a string of sites, each with a different adaptive value. Here, supergenes are either concatenation of neighboring genes (i.e., by physical location) or genes of the same ontology (by function). The merger would reduce false negatives due to low substitution numbers but at the risk of diluting true adaptive signal. We present the results based on the concatenations of neighboring genes in Table 1. In *Drosophila* and *Arabidopsis*, 200 and 500 supergenes are created, respectively. The results based on the merger by gene ontology are similar (see Table S1).

Our gene merger approach may create biases in the MK test, as pointed out before [3]. When the level of polymorphism is negatively correlated with the rate of non-synonymous divergence across loci, false positives would be common in the merger. Hence, we used the modified MK test to infer positive selection in merged genes [3]. In *Drosophila*, 112 of the 200 supergenes reject the MK test null hypothesis at the 5% level, and 36 of the 200 significantly deviate from the PAML null (Table 1). The two tests detect far more adaptive supergenes than individual genes: 56% (MK) and 18% (PAML). What is perplexing is that the overlap between the two sets is random (10.0% observed vs. the expected 10.1%), as if the two tests are completely uncorrelated. In *Arabidopsis*, 8.2% of the 500 supergenes pass the MK test at the 5% level, and 25.6% of supergenes reject the PAML null. The PAML test in *Arabidopsis* detects many more adaptive supergenes than the MK test, in the opposite direction of *Drosophila*. However, the overlap is also random, with 2.0% observed vis-à-vis the expected 2.1%. In both taxa, the two tests appear uncorrelated at the level of supergenes.

Because gene merger might dilute the adaptive signal by mixing a few adaptively evolving genes with many other non-adaptive genes, we examined the component genes within each adaptive supergene. In *Drosophila*, the 112 supergenes passing the MK test contain 3132 component genes (Table 1), among which 158 genes are significant when tested individually. Likewise, 60 out of 1040 component genes are identified by PAML. Between the two subsets of genes (3132 and 1040), 619 genes are common, and only three genes are significant by both tests. The 0.48% overlap of component genes is slightly higher than the expected 0.29%. The observations in *Arabidopsis* are given in the last row of Table 1. The overlap in component genes is also very low, at two of the 258 genes, or 0.78%. Clearly, the MK and PAML tests are uncorrelated by the standard statistical criteria. Comparable analyses using the PAML branch-site model (Table S2) yield results similar to those in Table 1.

#### 3. Identifying weakly adaptive genes with low stringency

We note in Fig. 2 that genes yielding a P value of 0.25 by either test may be moderately informative about positive selection. Therefore, when carrying out the MK and PAML tests simultaneously, we set the cutoff in each test at P < 0.224. By doing so, the expected overlap would be 0.224^2^ = 5% if the two tests are completely uncorrelated. The results of this relaxed stringency are given in Table 2.

**Table 2.** Proportion of adaptively evolving genes identified by two tests (P^2^ < 0.05, i.e. P < 0.224)

|  | MK | MK-PAML<br>overlap | PAML | Total |
| --- | --- | --- | --- | --- |
| *Drosophila* |  |  |  |  |
| No. of genes | 824 | 91<sup>d</sup> | 353 | 5425 |
| Expected overlap | / | 53.6 | / | / |
| Proportion of adaptive changes by MK<sup>a</sup> | 0.69 | 0.67 | 0.32 | 0.26 |
| No. of adaptive sites per gene by MK (A1) | 14.98 | 19.94 | 5.71 | 2.84 |
| No. of adaptive sites per gene by PAML (A2)<sup>b</sup> | 10.93 | 14.65 | 9.27 | 5.71 |
| No. of adaptive sites per gene by PAML (A2')<sup>c</sup> | 3.19 | 8.62 | 6.24 | 1.79 |
| *Arabidopsis* |  |  |  |  |
| No. of genes | 1014 | 119<sup>e</sup> | 1172 | 12975 |
| Expected overlap | / | 91.6 | / | / |
| Proportion of adaptive changes by MK | 0.69 | 0.67 | 0.06 | 0.04 |
| No. of adaptive sites per gene by MK (A1) | 19.36 | 28.98 | 1.97 | 0.84 |
| No. of adaptive sites per gene by PAML (A2) | 14.24 | 20.36 | 12.74 | 10.15 |
| No. of adaptive sites per gene by PAML (A2') | 3.59 | 11.51 | 8.13 | 2.44 |

<sup>a</sup> Proportion of adaptive changes is done using Shapiro et al. (2007)'s method of correction [3]
<sup>b</sup> A2 is based on the PAML-M2a model.
<sup>c</sup> A2' is based on the PAML-BEB model.
<sup>d</sup> $P < 10^{-7}$ by Fisher's exact test, given 53.6 as the expected value.
<sup>e</sup> $P < 0.002$ by Fisher's exact test, given 91.6 as the expected value.

The MK test identifies 824 and PAML 353 genes in *Drosophila*. These sets have 91 loci in common, whereas the expected overlap is 53.6 (P < 10^−7^, Fisher’s exact test). In *Arabidopsis*, the two tests yield 1014 and 1172 genes with an overlap of 119 genes, significantly higher than the expected number of 91.6 (P < 0.002, Fisher’s exact test). Hence, the joint call of adaptive genes accounts for 10.1% (119/1172) to 25.8% (91/353) of the loci identified by every single test. A gene identified by one test as adaptive has a 10% to 25% chance of being called adaptive by the other.

While the overlap between the two tests is at most modest, the performance of one test conditional on the pre-screen by the other indeed suggests some concordance. We first look at A1, the average number of adaptive sites per gene estimated using the MK test. A1 doubles from 2.84 to 5.71 when genes are pre-screened using PAML in *Drosophila* and increases from 14.98 to 19.94 in loci identified by both tests compared to just MK. The trend is even more pronounced in *Arabidopsis*: 0.84 to 1.97 and 19.36 to 28.98. Thus, the PAML screen can enhance the performance of the MK test.

The procedure is now applied in the reverse direction by pre-screening the genes with the MK test before subjecting them to the PAML test. The number of adaptive sites per gene can be calculated using two methods in PAML (A2 and A2’ in Table 2; Nielsen and Yang 1998 [40]; Zhang et al. 2005 [41]; see Supplementary Methods). Since the purpose is to compare PAML with MK, we use the A2 numbers, which are closer to A1 from the MK test. The qualitative conclusion, nevertheless, is not affected much by choice of model. The number of A2 sites increases from 5.71 to 10.93 after MK pre-screening in *Drosophila* (Table 2) and from 9.27 to 14.65 when focusing on the loci identified by both PAML and the MK test, compared to PAML alone. The same trend is observed in *Arabidopsis* (Table 2): an increase from 10.15 to 14.24 after MK test pre-screening and 12.74 for PAML only vs. 20.36 for genes identified by both tests. Again, a pre-screen by MK helps PAML performance. The results are similar when we use the PAML branch-site model rather than the site model (Table S3). It is clear that the MK and PAML tests are correlated, but the correlation is too weak to be of any practical use.

### PART II. The non-concordance between MK and PAML – Possible methodological reasons

In the first systematic comparisons of the MK and PAML tests on the same set of genes along the same phylogenetic branch, the detected adaptive signals are highly non-concordant. We first explore the possible technical reasons for the non-concordance.

#### 1. One test is right and the other is (nearly completely) wrong

Strong opinions have been expressed that either test is unreliable. This may also be the reason that few studies have used both tests to boost confidence, even when the data are amenable to both tests. However, as shown in Table 3, the two tests appear comparable in performance. When PAML is done on genes selected by the MK test, the subset of genes yields much stronger signal than the full set. This is also true when the MK test is done on PAML-selected genes. Since both tests have passed many prior simulations and applications [39,42–46] before becoming widely used, we now explore other explanations.

**Table 3.** MK and PAML tests on simulated sequence data.

| **Type of genes** | **Total number** | **PAML<sup>a</sup>**<br>**(% *false*)** | **MK<sup>b</sup>**<br>**(% *false*)** | **Overlap**<br>**(% *false*)** | **Concordance**<br>**(Overlap / MK)** |
| --- | --- | --- | --- | --- | --- |
| Individual genes<sup>c</sup> | 3000 | 1058 (5.5%) | 307 (17%) | 256 (0%) | 83.4% |
| Supergenes I<br>(constant selection)<sup>d</sup> | 300 | 297 (0%) | 169 (0.6%) | 168 (0%) | 99.4% |
| Supergenes II<br>(variable selection)<sup>e</sup> | 300 | 242 (0.8%) | 137 (2.9%) | 134 (0%) | 97.8% |

<sup>a</sup> Number of positive selected gene identified by the PAML site model test (Likelihood ratio test, $P < 0.05$ ).
<sup>b</sup> Number of positive selected gene identified by the MK test (Fisher's exact test, $P < 0.05$ ).
<sup>c</sup> There are 1000 positive selected genes and 2000 neutral genes in the simulated dataset.
<sup>d</sup> Every ten individual genes were merged into supergenes randomly.
<sup>e</sup> Every ten individual genes were merged into supergenes through a certain combination. A set of 60, 50, 30, 30, 30, 30, 20, 20, 10, 10 and 10 supergenes which respectively containing 0, 1, 2, 3, ..., 10 positive selected genes, was established to represent a case of variable selection.

#### 2. Both tests yield correct results, but for different aspects of positive selection

A common explanation for the non-concordance is that the two tests detect different aspects of positive selection. A more obvious scenario is the low power of the tests. We address the problem using supergenes. For example, the fraction of *Drosophila* supergenes yielding adaptive signals is 0.560 and 0.180, respectively, for MK and PAML (Table 1). The observed overlap at 0.100 is exactly the same as the overlap between random picks (0.101). Hence, high false-negative is not a correct explanation.

A set of more sophisticated scenarios is as follows: Since the evolution of DNA, sequencers proceed in a large space of parameters that vary in the strength, frequency, and mode of selection. It may be possible that some combinations of parameters may account for the discordance. For example, genes under strong selection, both positive and negative, may yield better signals in the MK test, while genes under weak selection may be more amenable to the PAML test. Most such conjectures can be tested by simulations. Nevertheless, given that the two tests were developed for the same purpose, the parameter sub-space yielding non-concordance might be very small and localized, if it can be found at all. We hence carried out extensive simulations that span a wider range of parameter values.

In simulating DNA sequence evolution, we allow all genes to harbor neutral and negatively selected sites. In a portion of these genes, a fraction of sites is further assumed to be driven by positive selection. Since an individual gene may not have a sufficiently large number of informative sites, we also bundle 20-30 randomly chosen genes into a supergene. Both false positives and false negatives are recorded. The test results on the simulated sequences are given in Fig. S2. Given the consistency, the condensed results are shown in Table 3.

In these simulations, PAML is more powerful than MK, although their relative power may be reversed under other conditions. The false positive rates are generally low. The number most relevant to this study is the concordance rate between the two tests (see Table 3). Because PAML is always more powerful in our simulations, the MK-detected genes are usually nested in the PAML-detected gene set. We therefore present the concordance rate as the number in the overlap (detected by both tests) divided by the number reported by the MK test. The concordance rate in our simulations is 83.4% for individual genes and close to 100% for supergenes. This high concordance rate thus presents a stark contrast with the results shown in Tables 1 and 2. The simulations show nearly full concordance between PAML and MK tests. In general, changing the parameters in the simulation would have substantial impacts on the detection rate of the two tests, but not their concordance. Therefore, the less sensitive method would detect a subset of genes reported by the more powerful method.

The efforts of Table 3 and Fig. S2 suggest that PAML and MK should generally be concordant. After all, they were developed for the same purpose. It is conceivable that the two tests might be less concordant than expected in some parts of the parameter space with the right combination of strength, frequency, and mode of selection. Such parameter combinations must be very rare as we could not find them in Fig. S2. Obviously, if genes that yield genuine incongruent signals between MK and PAML can be found, they must be driven by selection of a highly specific kind and would be most interesting. Nevertheless, instead of the continual search for the unusual, PART III below offers a much simpler explanation for the non-concordance.

### PART III. The biological reasons for the non-concordance – Fluctuating negative selection

In PART II, we reject the methodological reasons for the discordance between MK and PAML. We now propose a simple biological mechanism. As shown in Eq. (1’), most tests, including MK and PAML, have to assume constant *q* (the relative amount of negative selection) in the phylogeny of interest. Given the constancy, different tests may obtain *q* from various stages or lineages to achieve the same objective. We now test this fundamental assumption.

Eq. (2) shows that *q* can be estimated by Pa/Ps (∼ 1- *q*) from the polymorphism data within each species. Thus, a simple test of the constancy of *q* is to compare the Pa/Ps ratio in each species of interest. For example, between *A. thaliana* and *A. lyrata*, the Pa/Ps ratio is 0.152 and 0.248, and the Ka/Ks ratio is 0.184 (Table S4). Clearly, the strength of negative selection has changed in this short time span. In this case, the MK test would reach opposite conclusions depending on whether the polymorphism data used come from *A. thaliana* or *A. lyrata*. Obviously, if two MK tests do not agree, MK should not be expected to agree with the PAML test. How fluctuating negative selection would affect the PAML tests is more complicated since PAML is a collection of tests, each with a set of assumptions about how positive and negative selection operates [7,18,36,41]. How the PAML results are affected will be discussed in the Supplementary Notes.

We now use the polymorphism data (including the diploid genomes of single individuals) to investigate the variation in negative selection among extant species. The taxa are *Drosophila* (4 species), *Arabidopsis* (4 species or sub-species), primates (17 species), and birds (38 species). These data cover plants, invertebrates, and vertebrates.

#### 1. Drosophila

The four *Drosophila* species shown in Fig. 3a are taxa commonly used for probing adaptive molecular evolution [3,12,26,47]. Clearly, the selective constraint fluctuates wildly, even in this small group. Most notable is *D. sechellia*, which has a much higher Pa/Ps value than others. Among the rest, Pa/Ps at low frequency (<0.2) is higher in *D. melanogaster* than in *D. simulans* and *D. yakuba*, but above 0.2, the Pa/Ps values are similar (0.054-0.062, see Fig. 3a and Table S5).

**Figure 3.**
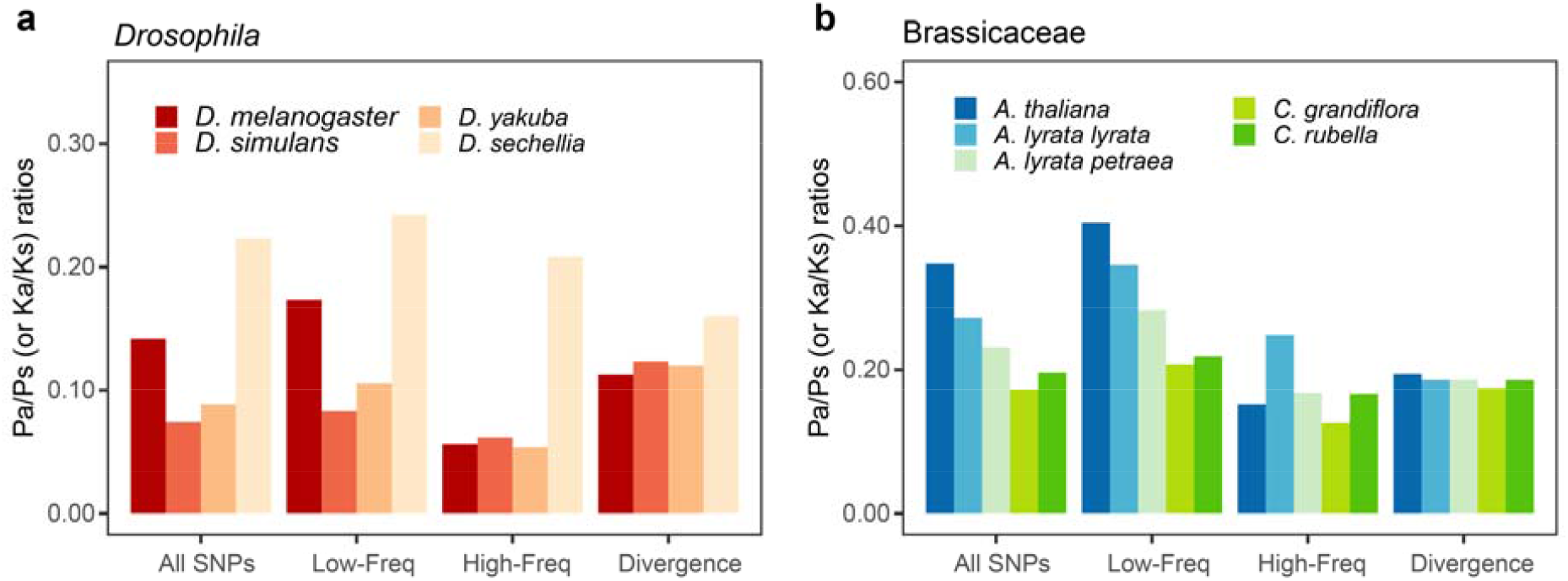
Pa/Ps ratios within species and Ka/Ks ratios between species. **a** and **b**, the Pa/Ps ratios grouped by frequency and Ka/Ks ratios in *Drosophila* and Brassicaceae. Variants with a frequency below 0.20 are grouped into Low-Freq bins, and other polymorphisms are grouped into High-Freq bins. The bars at Divergence bins represent lineage-specific Ka/Ks ratios.

The patterns raise some interesting issues. If one wishes to use the MK test to detect positive selection among the four species, one would compare the Pa/Ps ratio(s) with the Ka/Ks value between species. Fig. 3a shows that the lineage-specific Ka/Ks ratios are comparable, ranging between 0.113-0.160. Hence, the comparison between the interspecific Ka/Ks and the polymorphic Pa/Ps of *D. sechellia* would show no evidence of adaptive evolution. Among the remaining three species, the results are more nuanced. If all variants are used (as is commonly done), the conclusion for adaptive evolution would depend on the source of the polymorphism data – negative when using the data from *D. melanogaster* but positive when using the data from the other two species. Fay et al. (2002) proposed a cutoff of gene frequency depending on the polymorphism profile as the ones shown in Fig. 1c and 1d [38]. Many other procedures have since been introduced.

The main difficulty in detecting positive selection in *Drosophila* is that negative selection is not constant. Both MK and PAML may interpret the relaxation of negative selection as positive selection, albeit in different manners.

#### 2. Arabidopsis

*Arabidopsis thaliana* and its relatives are the main model organisms among plants, with high-quality reference genomes and polymorphism data [48,49]. The divergence of *A. thaliana* and *A. lyrata* is 11%, similar to the divergence between *D. melanogaster* and *D. simulans* (10.5%). However, negative selection in *Arabidopsis* fluctuates more wildly than in *Drosophila*. With the low-frequency SNPs (<0.2) removed, Pa/Ps is substantially higher in *A. lyrata* (*subsp. lyrata*) than in *A. thaliana* and *A. lyrata* (*subsp petraea*) (see Fig. 3b), whereas their lineage-specific Ka/Ks ratios are similar, ranging from 0.174 to 0.194. The results raise the question of the MK test again. When using the two taxa with the lower Pa/Ps (*A. thaliana* and *A. petraea*), one would conclude positive selection. But, using the ratios from *A. lyrata*, one would reach the opposite conclusion.

Compounding the issue, the polymorphism patterns are rather different among these species, and the cutoff to filter out the low-frequency variants should be different for each species (Fig. 3b and Table S4). *Arabidopsis* is, therefore, a typical example that the variation in the strength of negative selection, as manifested in Pa/Ps, is so large that the detection of positive selection based solely on DNA sequence data would be unreliable.

#### 3. Primates

For primates, we compile the data from 17 species (plus one subspecies) belonging to 6 genera. Since hominoids and old-world monkeys (OWMs) have diverged by < 6% in their DNA sequences, these two families are very close in molecular terms. This level of divergence would be suitable for detecting positive selection by both the MK and PAML test. (For a comparison, the two morphologically identical sibling species of *D. melanogaster* and *D. simulans* have diverged by more than 10%.)

The polymorphism and divergence data are presented Fig. 4a and Table S6. The polymorphism Pa/Ps ratio again varies considerably among species. The trend appears to be an increasing Pa/Ps ratio toward the human. With singleton and CpG sites removed, the polymorphism Pa/Ps ratio in primates decreases in the following order: human and bonobo (0.400-0.384), chimpanzee and gorilla (0.353-0.295), orangutan (0.298-0.282) and macaque monkeys (0.245-0.237) (see Table S6 and Table S7). The snub-nosed monkey (0.349-0.332), an old-world monkey, is the only group that deviates from the general trend.

**Figure 4.**
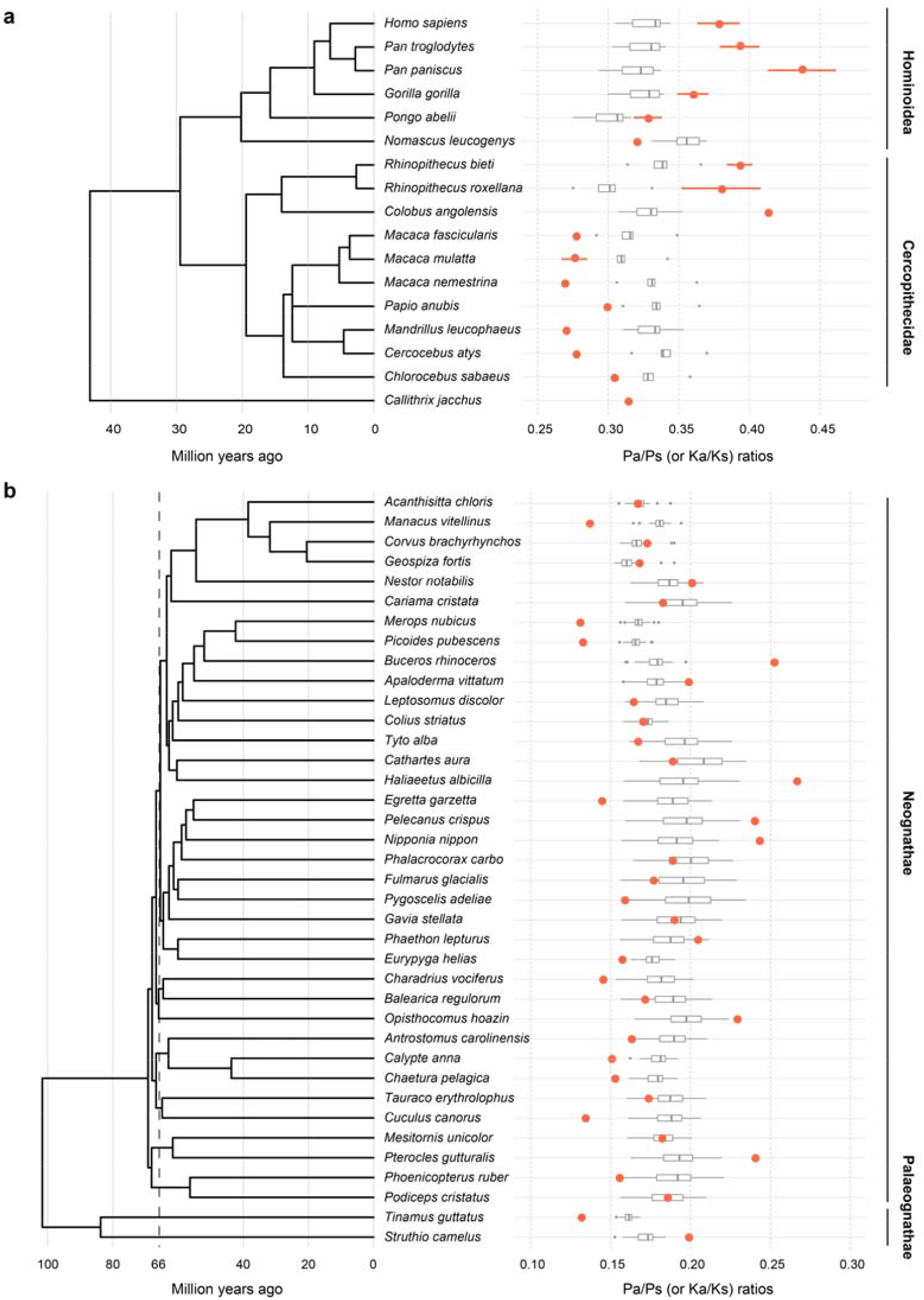
Pa/Ps ratios within species and Ka/Ks ratios between species. **a** and **b**, Pa/Ps (or Ka/Ks) ratios in primates and birds. The red points show the heterozygous Pa/Ps ratios. Since eight species of primates have genomic data of more than five individuals, their means and standard deviations of heterozygous Pa/Ps ratios are displayed as red points and attached error bars. Each grey boxplot represents the distribution of Ka/Ks ratios between this species and other species on the phylogenetic tree. Note that the Ka/Ks ratios between species in the same family are excluded in panel **a**.

The trend of a larger Pa/Ps ratio toward apes and the human casts doubts on the inferences of adaptive evolution in the genomic sequences of primates. Note that Ka/Ks > Pa/Ps is often assumed to be the hallmark of positive selection. The Ka/Ks ratio for each lineage, say human (any ape species), is the comparison with every OWM species, thus yielding a distribution as shown. Likewise, each OWM species is compared with every ape. If we compare the human and any OWM species, the Ka/Ks value is 0.305-0.344. Since the Pa/Ps is human is about 0.382, there is no evidence of adaptive evolution between human and OWMs (see Table S6 and Table S7). However, since Pa/Ps ranges between 0.269 and 0.304 among OWMs, one would conclude positive selection between the same two taxa.

The contradiction just stated is very general when one compares any ape species with any OWM species. In Fig. 4a, the top half (including all apes, *Rhinopithecus* and *Colobus*) shows that the Pa/Ps ratios (red dots) generally fall in the higher range of 0.320-0.413 while the 7 OWM species in 5 genera of the lower half have Pa/Ps between 0.269 and 0.304. Importantly, the Ka/Ks ratio generally falls between the two sets of Pa/Ps values. In short, the assumption of constant negative selection is violated between apes and OWMs, thus precluding the inference of adaptive evolution in genomic sequences between them.

#### 4. Birds

For birds, we analyze the genomic data of 38 species from 30 orders. The general trend is the same as those of the three taxa shown in Fig. 3 and Fig. 4a. In Fig. 4b, the Pa/Ps ratios are scattered between 0.131 and 0.266 among all bird species with a mean of 0.179 and a standard deviation of 0.035 (see Table S8). In contrast, the Ka/Ks ratio falls in a relatively small range of 0.152-0.235. Again, the variation in the strength of negative selection in birds, as seen in the Pa/Ps value relative to the interspecific divergence in Ka/Ks, is far too large to permit the analysis of positive selection based solely on the genomic sequences.

## Discussion

A central task of molecular evolution studies is to quantify the amount of positive selection acting on the genomes. The identification of genes under positive selection will then permit further functional studies. The ultimate goal is to connect adaptive evolution to its mechanistic bases. In the neutral theory that has dominated the field in the last 50 years, the genomic sequences, driven mainly by negative selection and genetic drift, carry little signals of adaptive evolution [50–54].

For two decades, the search for signals of adaptive sequence evolution appears to have strongly refuted the neutral theory. However, the studies have been done under a fundamental constraint in the analysis of genomic evolution between species. The constraint is that negative selection (*q* of Eq 1’) has to be a constant pressure if one wishes to extract information about positive selection from the sequences. If *q* is a constant, it, in principle, can be estimated from any lineage and at any stage of evolution. Thus, although MK and PAML differ in how they estimate *q*, the results are expected to converge. However, it is surprising how weakly the two methods converge, as shown in Tables 1 and 2.

In this current report, negative selection is found to deviate strongly from the assumed constancy in all taxa analyzed. Hence, the reported adaptive evolution in DNA sequences in the last two decades by either MK or PAML could be seriously confounded by the fluctuation in negative selection. Given that positive and negative selection may both fluctuate in time and between lineages, could the extraction of information on adaptive evolution from genomic sequences, in fact, be theoretically untenable?

In light of the current report, we now outline a path forward that is elaborated in the companion study [55]. In taking the new path, we are concerned with measuring the fluctuation in negative selection without addressing the many factors underlying the fluctuation. These factors may include environment, genetic background, population size, and so on, and are briefly commented on in the Supplementary Notes. The central idea of the path forward is to analyze genomic sequences in two stages. In the first stage, a complete map of negative selection, including the frequency and strength of negative selection and the variation in time and across lineages, will be worked out for the phylogeny of interest. This is feasible if the number of deleterious mutations is much larger than that of beneficial ones, i.e., *q* >> *p*. In the second stage, the inference of positive selection will then be built on this detailed map of negative selection. It is important to note that prior efforts have been cursory in stage I by assuming constancy of negative selection.

In the first stage, it is necessary to track the changes in negative selection in each lineage through time. This can be done indirectly by estimating the effective population size at each evolutionary time interval using the PSMC method [56] or the step-ladder method [57]. It is the basic population genetic principle that the larger the effective population size (*N*_*e*_), the more effective selection would be. Hence, the Pa/Ps ratio would be lower due to stronger negative selection when *N*_*e*_ becomes larger. With the knowledge of *N*_*e*_ changes, it is often (but not always) possible to know the changes in negative selection. In Chen et al. (2020), the indirect inferences of changes in negative selection through time as a function of *Ne* changes have been done in several species of primates [55]. Most interesting, by using PSMC, one could take the direct approach to negative selection by calculating Pa/Ps for each time interval. In short, a detailed map of changes in negative selection may be feasible although the methodologies are still incompletely developed.

Finally, the strengths of positive selection and negative selection are strongly correlated (partly because both are functions of *N*_*e*_). For example, large-step mutations, when measured by the physicochemical distances between amino acids, are both more deleterious and more beneficial than small-step mutations [21,22]. Therefore, the working of negative selection would be informative about the operation of positive selection and vice versa. In conclusion, to understand adaptive evolution at the DNA level, we must have a complete understanding of negative selection first and in the same context.

## Materials and Methods

The DNA sequence data of *Drosophila*, Brassicaceae, primates and birds were obtained from the public database. The detailed information of sequence database and screening process for each category is given in the Supplementary Methods.

We used an approximate method to estimate Ka/Ks ratio with an improved Nei-Gojobori model [11,20,58] Simple methods for estimating the numbers Simple methods for estimating the numbers. The values of Pa and Ps were observed from polymorphism data using the same method. To avoid the confounding effect of negative selection on the MK test, we only used common mutations with derived allele frequencies larger than 0.2, as was done previously [3,59]. We used both the site model and the branch-site model in PAML. The detailed methods of MK and PAML test, the simulations of coding sequence evolution, and the supergene construction are also given in the Supplementary Methods.

## Acknowledgements

We thank Daniel Hartl, Ziheng Yang, Adam Eyre-Walker, Dan Graur, Justin Fay, Bruce Rannala, and Matthew Hahn for their helpful discussions. This study was supported by the National Natural Science Foundation of China (31971540 and 31830005), the National Key Research and Development Plan (2017FY100705), the Guangdong Basic and Applied Basic Research Foundation (2019A1515010752), the Science and Technology Project of Guangzhou (202102020483), and the Innovation Group Project of Southern Marine Science and Engineering Guangdong Laboratory (Zhuhai) (No. 311021006).

## References

1. Fay JC, Wu C-I. Sequence divergence, functional constraint and selection in protein evolution. Annu Rev Genomics Hum Genet 2003;4:213–35.

2. Smith NGC, Eyre-Walker A. Adaptive protein evolution in Drosophila. Nature 2002;415:1022–4.

3. Shapiro JA, Huang W, Zhang C et al. Adaptive genic evolution in the Drosophila genomes. Proc Natl Acad Sci U S A 2007;104:2271–6.

4. Wang H-Y, Chen Y, Tong D et al. Is the evolution in tumors Darwinian or non-Darwinian? Natl Sci Rev 2018;5:15–7.

5. Wen H, Wang H-Y, He X et al. On the low reproducibility of cancer studies. Natl Sci Rev 2018;5:619– 24.

6. Boyko AR, Williamson SH, Indap AR et al. Assessing the evolutionary impact of amino acid mutations in the human genome. PLoS Genet 2008;4:e1000083.

7. Yang Z, Swanson WJ. Codon-substitution models to detect adaptive evolution that account for heterogeneous selective pressures among site classes. Mol Biol Evol 2002;19:49–57.

8. Yang Z, Wong WSW, Nielsen R. Bayes empirical Bayes inference of amino acid sites under positive selection. Mol Biol Evol 2005;22:1107–18.

9. Li H, Stephan W. Inferring the demographic history and rate of adaptive substitution in Drosophila. PLOS Genet 2005;2:e166.

10. Tajima F. Statistical method for testing the neutral mutation hypothesis by DNA polymorphism. Genetics 1989;123:585–95.

11. Nei M, Gojobori T. Simple methods for estimating the numbers of synonymous and nonsynonymous nucleotide substitutions. Mol Biol Evol 1986;3:418–26.

12. McDonald J, Kreitman M. Adaptative protein evolution at the Adh locus in Drosophila. Nature 1991;354:293–5.

13. Fay JC, Wu C-I. Hitchhiking under positive Darwinian selection. Genetics 2000;155:1405–13.

14. Yang Z, Bielawski JP. Statistical methods for detecting molecular adaptation. Trends Ecol Evol 2000;15:496–503.

15. Sabeti PC, Reich DE, Higgins JM et al. Detecting recent positive selection in the human genome from haplotype structure. Nature 2002;419:832–7.

16. Sawyer SA, Hartl DL. Population genetics of polymorphism and divergence. Genetics 1992;132:1161–76.

17. Li W-H, Wu C-I, Luo C-C. A new method for estimating synonymous and nonsynonymous rates of nucleotide substitution considering the relative likelihood of nucleotide and codon changes. Mol Biol Evol 1985;2:150–74.

18. Yang Z, Swanson WJ, Vacquier VD. Maximum-likelihood analysis of molecular adaptation in abalone sperm lysin reveals variable selective pressures among lineages and sites. Mol Biol Evol 2000;17:1446–55.

19. Fay JC, Wyckoff GJ, Wu C-I. Positive and negative selection on the human genome. Genetics 2001;158:1227–34.

20. Tang H, Wyckoff GJ, Lu J et al. A universal evolutionary index for amino acid changes. Mol Biol Evol 2004;21:1548–56.

21. Chen Q, He Z, Lan A et al. Molecular evolution in large steps-codon substitutions under positive selection. Mol Biol Evol 2019;36:1862–73.

22. Chen Q, Lan A, Shen X et al. Molecular evolution in small steps under prevailing negative selection: a nearly universal rule of codon substitution. Genome Biol Evol 2019;11:2702–12.

23. Nielsen R, Yang Z. Estimating synonymous and nonsynonymous substitution rates under realistic evolutionary models. Mol Biol Evol 2000;17:32–43.

24. Li W-H. Molecular Evolution. Sunderland (MA): Sinauer Associates, 1997.

25. Chen B, Shi Z, Chen Q et al. Tumorigenesis as the paradigm of quasi-neutral molecular evolution. Mol Biol Evol 2019;36:1430–41.

26. Clark AG, Eisen MB, Smith DR et al. Evolution of genes and genomes on the Drosophila phylogeny. Nature 2007;450:203–18.

27. Nielsen R, Bustamante CD, Clark AG et al. A scan for positively selected genes in the genomes of humans and chimpanzees. PLoS Biol 2005;3:0976–85.

28. Yang Z. PAML: a program package for phylogenetic analysis by maximum likelihood. Bioinformatics 1997;13:555–6.

29. Yang Z. PAML 4: phylogenetic analysis by maximum likelihood. Mol Biol Evol 2007;24:1586–91.

30. Smith MD, Wertheim JO, Weaver S et al. Less is more: An adaptive branch-site random effects model for efficient detection of episodic diversifying selection. Mol Biol Evol 2015;32:1342–53.

31. Kosakovsky Pond SL, Frost SDW, Muse S V. HyPhy: Hypothesis testing using phylogenies. Bioinformatics 2005;21:676–9.

32. Kosakovsky Pond SL, Poon AFY, Velazquez R et al. HyPhy 2.5 - A customizable platform for evolutionary hypothesis testing using phylogenies. Mol Biol Evol 2020;37:295–9.

33. Gillespie JH. Population Genetics: A Concise Guide. 2nd ed. Baltimore: The John Hopkins University Press, 2004.

34. Ohta T, Gillespie JH. Development of neutral nearly neutral theories. Theor Popul Biol 1996;49:128–42.

35. Kimura M. Some problems of stochastic processes in genetics. Ann Math Stat 1957;28:882–901.

36. Yang Z, Nielsen R. Codon-substitution models for detecting molecular adaptation at individual sites along specific lineages. Mol Biol Evol 2002;19:908–17.

37. Portik DM, Leaché AD, Rivera D et al. Evaluating mechanisms of diversification in a Guineo-Congolian tropical forest frog using demographic model selection. Mol Ecol 2017;26:5245–63.

38. Fay JC, Wyckoff GJ, Wu C-I. Testing the neutral theory of molecular evolution with genomic data from Drosophila. Nature 2002;415:1024–6.

39. Messer PW, Petrov DA. Frequent adaptation and the McDonald-Kreitman test. Proc Natl Acad Sci U S A 2013;110:8615–20.

40. Nielsen R, Yang Z. Likelihood models for detecting positively selected amino acid sites and applications to the HIV-1 envelope gene. Genetics 1998;148:929–36.

41. Zhang J, Nielsen R, Yang Z. Evaluation of an improved branch-site likelihood method for detecting positive selection at the molecular level. Mol Biol Evol 2005;22:2472–9.

42. Fu W, Gittelman RM, Bamshad MJ et al. Characteristics of neutral and deleterious protein-coding variation among individuals and populations. Am J Hum Genet 2014;95:421–36.

43. Anisimova M, Nielsen R, Yang Z. Effect of recombination on the accuracy of the likelihood method for detecting positive selection at amino acid sites. Genetics 2003;164:1229–36.

44. Wong WSW, Yang Z, Goldman N et al. Accuracy and power of statistical methods for detecting adaptive evolution in protein coding sequences and for identifying positively selected sites. Genetics 2004;168:1041–51.

45. Suzuki Y, Nei M. Simulation study of the reliability and robustness of the statistical methods for detecting positive selection at single amino acid sites. Mol Biol Evol 2002;19:1865–9.

46. Eyre-Walker A. Changing effective population size and the McDonald-Kreitman test. Genetics 2002;162:2017–24.

47. Halligan DL, Oliver F, Eyre-Walker A et al. Evidence for pervasive adaptive protein evolution in wild mice. PLoS Genet 2010;6:e1000825.

48. Alonso-Blanco C, Andrade J, Becker C et al. 1,135 genomes reveal the global pattern of polymorphism in Arabidopsis thaliana. Cell 2016;166:481–91.

49. Weigel D, Mott R. The 1001 Genomes Project for Arabidopsis thaliana. Genome Biol 2009;10:107.

50. Bustamante CD, Fledel-Alon A, Williamson S et al. Natural selection on protein-coding genes in the human genome. Nature 2005;437:1153–7.

51. Biswas S, Akey JM. Genomic insights into positive selection. Trends Genet 2006;22:437–46.

52. Welch JJ. Estimating the genomewide rate of adaptive protein evolution in Drosophila. Genetics 2006;173:821–37.

53. Gayà-Vidal M, Albà MM. Uncovering adaptive evolution in the human lineage. BMC Genomics 2014;15:599.

54. Luisi P, Alvarez-Ponce D, Pybus M et al. Recent positive selection has acted on genes encoding proteins with more interactions within the whole human interactome. Genome Biol Evol 2015;7:1141–54.

55. Chen Q, He Z, Feng X et al. Two decades of suspect evidence for adaptive DNA-sequence evolution – Less negative selection misconstrued as positive selection. bioRxiv 2020, DOI: 10.1101/2020.04.21.049973.

56. Li H, Durbin R. Inference of human population history from individual whole-genome sequences. Nature 2011;475:493–6.

57. Liu X, Fu YX. Exploring population size changes using SNP frequency spectra. Nat Genet 2015;47:555–9.

58. Ina Y. New methods for estimating the numbers of synonymous and nonsynonymous substitutions. J Mol Evol 1995;40:190–226.

59. Akashi H. Within- and between-species DNA sequence variation and the “footprint” of natural selection. Gene 1999;238:39–51.

