## Supplementary information for "Two decades of suspect evidence for adaptive molecular evolution – Negative selection confounding positive selection signals"

**This file includes:**

Supplementary Methods

Supplementary Notes

Supplementary References

Figures S1-S4

Tables S1-S6

**Supplementary Methods**

***DNA sequence data***

1) Genus *Drosophila*

Coding sequences (CDS and their translated proteins) and orthologs information of *D. melanogaster* were downloaded from FlyBase (http://www.flybase.org) [60,61]. We used the FB2019_03 gene models. Protein sequences were aligned using default parameters in MUSCLE v3.8.31 [62]. We then aligned the CDS of orthologous genes with the help of aligned protein sequences using PAL2NAL v14 [63]. We collected 8560 alignments of five species (*D. melanogaster, D. simulans, D. sechellia, D. yakuba,* and *D. erecta*, Fig. 1a) spanning 11 million years of evolution (see Fig. S3) [26,64,65].

We downloaded 99 *D. melanogaster* genomes from the *Drosophila* Population Genomics Project 2 (DPGP2) website [66]. *Drosophila simulan*s polymorphism data from 170 inbred individuals were obtained from Signor et al. (2018) [67]. Sequence data for 20 *D. yakuba* inbred lines are from Rogers et al. (2014) [68]. *D. sechellia* data are from 41 outbred lines described in Schrider et al. (2018) [69]. We aligned reads from each line using the BWA_MEM algorithm of BWA 0.7.17 to the appropriate reference sequences (Table S9) [70]. We then used SAMtools 1.9 to sort, index alignments, and pick properly paired alignments [71]. We used Picard (https://github.com/broadinstitute/picard) to mark and remove duplicates and performed local realignment with GATK's RealignerTargetCreator and IndelRealigner tools (version 3.7) [72]. We used BCFtools mpileup with the option "-Q 20 -q 20 -a DP, AD" and bcftools call with the option "-cvO" for variant calling [73]. Multi-allelic sites and indels were excluded from analyses using VCFtools v0.1.17 [74] option "--remove-indels --min-alleles 2 --max-alleles 2".

Genes with high divergence rates, apparently caused by misalignment, were discarded. Only genes with more than 40 codons and ten samples of polymorphism data were used. The final dataset contains 5245 genes with an average of 50 samples of polymorphism data.

2) Family Brassicaceae

The DNA sequences of *A. thaliana*, *A. lyrata*, *Capsella grandiflora*, *C. rubella*, and *Boechera stricta* were obtained from the PhytozomeV12 database [75–79]. Pairwise orthologs for the five species were inferred using the InParanoid algorithm [80]. Only genes with a single 1:1 ortholog for each species were used. We began with 14953 alignments of the five species and then did the alignment and filtered the data as we did for *Drosophila*.

We used *A. thaliana* genome-wide polymorphism data sets from the 1001 Genomes Project [48,49]. Briefly, 1135 wild inbred lines were used for this study. Raw *Arabidopsis lyrata* sequence data were downloaded from NCBI, and genotypes were called followed the protocol of Mattila et al. (2017) [81]. We obtained sequence data from 8 *Capsella grandiflora* samples from Josephs et al. (2015) [82] and 12 *C. rubella* samples from Ågren et al. (2014) [83]. We used BWA 0.7.17 [70], SAMtools 1.9 [71], and GATK 3.7 [72] to generate genotype calls as described above for *Drosophila*. Only genes with more than 300 *A. thaliana* samples of polymorphism data were collected. The final dataset consists of 12975 orthologous genes.

3) Order Primates

In the order Primates, 16 Catarrhini species have genome assemblies in NCBI (The National Center for Biotechnology Information) [84–96]. We used the common marmoset genome data as the outgroup [97]. A summary of assemblies and annotations of these 17 species is found in Table S9. The phylogenetic tree was downloaded from the Time Tree website (http://www.timetree.org/). Using the software OrthoFinder v2.2.7 [98], optimized with Diamond [99], we identified 7538 autosomal *Homo sapiens*-centric orthologous gene sets of 17 primate species. CDS and protein alignments were obtained using MUSCLE v3.8.31 and PAL2NAL v14 [62,63]. Because mammals exhibit hyper-neutral CpG mutation rates [100,101], we masked codons with CpGs to TpG/CpAs changes.

We downloaded genotypes of Han Chinese and Yoruba individuals as VCF files from the 1000 Genomes Project website [102,103]. We obtained genotypes of bonobos, chimpanzees, gorillas, and orangutans from Prado-Martinez et al. (2013) [104] and the genome-wide polymorphism data set of four subspecies of Chinese rhesus macaques *Macaca mnulatta* from Liu et al. (2018) [105]. We obtained sequence data from 18 snub-nosed monkeys *Rhinopithecus. roxellana* samples and 20 *R. bieti* samples from Yu et al. (2016) [96]. Finally, we downloaded eight old world monkeys and one marmoset genome (*Callithrix jacchus*) raw reads from NCBI (listed in Table S9). We again used BWA0.7.17 [70], SAMtools 1.9 [71], and GATK 3.7 [72] to generate genotype calls from the raw read data sets.

4) Class Aves

The CDS and translated sequences of protein-coding genes of 38 birds were obtained from the Avian Phylogenomic Project website [106–112]. We used OrthoFinder v2.27 with the aid of Diamond to find 8295 orthologs [98,99]. The MUSCLE-PANLNAL pipeline described above was used to align protein-coding sequences. Raw reads for genome assembly were downloaded from NCBI to obtain polymorphisms from single individuals. A summary of genome assemblies and raw reads could be found in Table S9. The phylogenetic tree of Fig. 4b was modified from the highly resolved total evidence nucleotide tree inferred with ExaML by Jarvis et al. (2014) [108]. We used BWA 0.7.17, SAMtools1.9, and GATK 3.7 to generate genotype calls [70–73]. The pipelines are described in more detail in the *Drosophila* section.

***Supergene construction***

To overcome statistical limitations, we created artificial supergenes by merging genes into longer sequences. We used two concatenation approaches: by physical location and by ontology. The first method involved merging 20 to 30 nearby genes residing on the same chromosome. This resulted in 200 *Drosophila* and 500 *Arabidopsis* supergenes. To apply the ontology approach, we first identified GO (gene ontology) term(s) for each gene. To ensure that every gene was present in only one supergene, we sorted GO terms by the number of genes they comprised and checked the component genes in each supergene. If a gene was previously included in a set, it was not merged again. GO terms with fewer than eight genes in *Drosophila* and ten in *Arabidopsis* were discarded. The final set comprised 184 *Drosophila* and 454 *Arabidopsis* supergenes.

***Simulations of coding sequence evolution***

For each simulated gene, we used software SLiM3 (an evolutionary simulation framework) [113], which supports the Wright-Fisher model and nucleotide-based model, to do forward simulations to obtain protein-coding sequences of five species (*SP1*, *SP2*, *SP3*, *SP4*, and *SP5*). The divergence time is the same as the phylogenetic tree of *Drosophila* (see Fig. S3). We set constant population size $N$ = 100000 for each species, mutation rate $\mu = 1\times10^{-9}$/site/year, $\kappa = 2$ in Kimura (1980) mutation model [114]. In order to simplify the simulation process, we set one generation per year for each species. Each gene consists of 2000 codons. We made all synonymous mutations neutral and nonsense mutations lethal. And the distribution of the fitness effect of non-synonymous mutations is set to a combination of parameters of different values and proportions (see Fig. S2). There are 1000 positive selected genes and 2000 neutral genes in the simulated dataset. For the 1000 simulated positive selected gene, as used in Table 3, there are 10% ($p$) beneficial mutations ($Ns = 5$), 85% ($q$) deleterious mutations ($Ns = -50$) and 5% ($1 - p - q$) neutral mutations. For the 2000 neutral genes, there are no beneficial mutations ($p = 0$), $q$ (85%, $Ns = -50$) deleterious mutation and $1 - p - q$ (15%) neutral mutations. To improve simulation performance, we rescaled population size $N$ by a factor 100 under the guidance of SLiM manual [115]. We then rescaled $\mu$ and $s$ such that the product $N\mu$ and $Ns$ remain the same. After simulation, we sampled one sequence of each species to do PAML test. And we sampled sequences of 100 individuals in *SP1* and one individual in *SP2* to do MK test.

We merged the 3000 genes used in Table 3 into 300 supergenes. We first merged every ten genes into a supergene randomly. There are 3, 38, 51, 75, 66, 43, 14, 8 and 2 supergenes containing 0, 1, 2, 3, 4, 5, 6, 7 and 8 positive selected genes, respectively. We also merged every ten genes into a supergene through a certain combination. In this case, there are 60, 50, 30, 30, 30, 30, 20, 20, 10, 10 and 10 supergenes containing 0, 1, 2, 3, …, 10 positive selected genes, respectively. In the MK test, the corrections suggested by Shapiro et al. (2007) are not implemented because the concordance rate remains virtually identical with or without the corrections [3].

***Computation of Ka/Ks ratios and Pa/Ps ratios***

We used an approximate method to estimate Ka/Ks ratio with an improved Nei-Gojobori model [11,20,58]. We estimated transition to transversion ratio ($\kappa/2$ in Nielsen and Yang 1998 [40]) from the fourfold degenerate sites. We then used this ratio to count the expected number of synonymous and non-synonymous mutations in each codon. Next, we counted the number of synonymous and non-synonymous substitutions. When more than two differences exist between any two codons compared, we used the method described in Nei and Gojobori (1986) [11]. Finally, we used the Jukes-Cantor (1969) [116] method to correct for multiple substitutions and obtain substitution rate estimates (Ka and Ks).

Because some genome assemblies used for polymorphisms calls have been updated, we changed the genome versions of polymorphism data of some species. For example, we downloaded the LiftOver file from UCSC and converted the genome coordinates of DPGP2 variants from dmel5 to dmel6 using CrossMap [117] (see Table S9). We then used SnpEff [118] to annotate synonymous and non-synonymous changes for polymorphism based on the reference assembly. For *Drosophila* and *Arabidopsis* data, we used the free ratios model (model = 1, NSsites = 0, fix_omega = 0) of CODEML PAML module [28,29] to infer ancestral sequences at each node on the known phylogenetic trees (see Fig. S3 and Fig. S4). The Ka/Ks ratios of each branch were then estimated using the method mentioned above. The identical allele corresponding to the ancestral branch was set as the inherited allele, and the alternative was derived. Thus, we were able to estimate the derived allele frequency. The computation of Pa/Ps ratios is similar to the method for the Ka/Ks ratio, which is described above. We obtained Pa and Ps directly from observed polymorphism data and expected mutation counts without multiple hits correction.

***The McDonald-Kreitman (MK) test***

In Fig. 1c-1d, Pa/Ps ratio is given for the number of polymorphic changes at a defined frequency range. The Pa/Ps ratio becomes lower when the mutant frequency becomes higher. Apparently, the

Pa/Ps ratio at the low-frequency range is boosted by deleterious mutations that have not been removed by negative selection. To avoid the confounding effect of negative selection on the MK test, we only used common mutations with derived allele frequencies larger than 0.2, as was done previously [3,59]. Note that the Pa/Ps ratio reaches a steady level at around 0.2 in Fig. 1c-1d.

Now, we let PolyA and PolyS designate the total number of common polymorphic mutations per gene (frequency > 0.2) in the MK test. The corresponding numbers of changes between species are designated DivA and DivS. These four numbers are gathered in a 2×2 contingency table. Fractions of amino acid substitutions which are adaptive can be estimated as 1 - (PolyA / PolyS) / (DivA / DivS). We used Fisher's exact test on 2×2 contingency tables to estimate statistical significance. Shapiro et al. (2007) pointed out the possibility of false positives when the MK test is applied across genes and proposed a procedure to correct the bias [3]. Hence, we used it in the calculations.

***The PAML test***

We used both the site model and the branch-site model in PAML. The site model, allowing the ω ratio ($d_N/d_S$, or Ka/Ks) to vary among sites (codons or amino acids in protein), detected positive selection across the five chosen species. A likelihood ratio test (LRT) was used to compare the alternative model M2a (selection model allowing an additional category of positively selected sites with $\omega > 1$ by setting: model = 0, NSsites = 2, fix_omega = 0, omega = 2) with the null model M1a (neutral model allowing only two categories of sites with $\omega < 1$ and $\omega = 1$ by setting: model = 0, NSsites = 1, fix_omega = 0, omega = 2). Significance was determined using the chi-squared test (df = 2).

The branch-site model, allowing $\omega$ to vary both among sites and across lineages, was used to detect positive selection along specified branches. We compared the likelihood of the alternative model A (positive selection, model = 2, NSsites = 2, fix_omega = 0), to the null model A1 (model = 2, NSsites = 2, fix_omega = 1, omega = 1). *D. melanogaster* and *A. thaliana* were designated as the foreground branches for the test. Significance was calculated using LRT as above. The site model results are presented in the text, and the branch-site results are given in Supplementary Notes.

In the analysis of both models, we also employed Bayes empirical Bayes (BEB) [8] estimates, which are available for calculating the posterior probabilities for site classes and can be used to identify sites under positive selection if the likelihood ratio test is significant.

### Supplementary Notes

#### *On selection signals in interspecific (divergence) vs. intraspecific (polymorphism) comparisons*

The detection of positive selection largely falls into two broad classes [10–14]. One class attempts to detect positive selection that operates within populations [10,13,15]. The other focuses on positive selection that operates in the longer term, i.e., the divergence between species [16–18]. In this study, we focus on the between-species tests. Technically, there is indeed more room to develop new methods to detect positive selection within species. This is particularly relevant to human population genetic studies which focus almost entirely on polymorphisms. We have contributed to the detection of within-population selection as done in Ref. [13,119–121].

Now, we wish to answer the question, "what might be the differences between positive selection within vs. between species?" While the question is only tangential to our study, it is conceptually much more satisfying to detect selection between species. In the case of primates, the two papers attempt to detect positive selection of the last 6 Myrs (million years) between human and chimpanzee and in the last 35 Myrs between humans and macaque. In contrast, the polymorphism data may inform about selection of the last 0.3 Myrs.

A lesson from *Drosophila* is most instructive. Numerous studies have identified many quantitative trait loci (QTLs) for the variation in bristle number in *D. melanogaster*. The possible selective advantages of these QTLs have been the focus of further analyses. There is, however, a curious dilemma. The bristle number is, in fact, quite stable between *Drosophila* species, even distantly related ones. The contrast may mean two things. First, the signals detected from the polymorphism data are noises. Second, the signals are truly adaptive ones, and selection is mainly fine-tuning near the adaptive equilibrium. In short, the long-term adaptive evolution is different in nature than short term signals detected in polymorphism. Population geneticists are aware of the dilemma.

It is essential that we analyze positive selection between species independent of works done on adaptive signals within species. They are two complementary, rather than redundant, approaches. We believe that MK and PAML are important for this reason.

#### *Previous studies that employed MK and PAML tests*

Some previous studies did use both MK test and PAML, but their purposes were varied. They can be roughly grouped into several categories. The first category used both of their results to make a conclusion but did not discuss too much about their inconsistencies. For instance, Luisi et al. (2015) studied how positive selection effects across parts of molecular networks using comparative genomics and population genetics approaches on human and related species' genomes [54]. This article concluded that the relationship between centrality and the impact of adaptive evolution highly depends on the mode of positive selection and/or the evolutionary time scale. Feldmeyer et al. (2015) used three methods (MK test, PAML branch model, and TreeSAAP) to identify positively selected genes involved in species divergence on orthologous mt-genome transcripts of four closely related European Radix species [122]. In total, 134 genes were identified as positively selected by at least one of the three methods. Of these, 116 were identified by a single method only, and 5 genes by all three methods. Wong et al. (2007) proposed that seminal fluid proteases are likely targets of selection due to their demonstrated or potential roles in between-sex interactions and immune processes. The gene pool is limited to five predicted protease-encoding *Acp* loci to test this hypothesis, and several within- and between-species sequence methods were used, including the MK test and PAML [123]. One gene was identified by the MK test whereas PAML found another two. Biswas et al. described the identification of targets of adaptive evolution [51]. They reviewed previous studies using different methods to detect positive selection based on polymorphisms within species, as well as polymorphism within and divergence between species. Some inconsistencies in these results have been observed but this paper offers three simple explanations: 1) different studies are probably detecting different selective events; 2) even for tests that should detect similar types of selective events, low statistical power further decreases the probability of overlap; 3) most studies report only the most significant results.

The second category uses one method to detect adaptive evolution and adopt the idea from the other as a supplement. For example, Nielsen et al. (2005) compared 13731 annotated genes from humans to their chimpanzee orthologs to identify genes that show evidence of positive selection [27]. They identified 50 genes with the highest likelihood ratio. PAML was used to validate those genes if the elevated $d_N/d_S$ was in the human lineage, chimpanzee lineage, or both lineages. Population genetic analysis of non-synonymous polymorphisms of those 50 genes showed evidence for an excess of high frequency alleles, providing additional support for positive selection. Gayà-Vidal and Albà (2014) used the MK test for estimating the amount of positive selection in the human lineage [53]. It considered several factors that could have an impact on the fraction of non-synonymous substitutions ($\alpha$): 1) high derived allele frequency (up to 60%) will avoid the impact of slightly deleterious mutations and thus elevate $\alpha$; 2) Human accelerated genes ($d_N/d_S$ accelerate in human branch) and human BS test genes (genes detected by PAML branch-site model) would elevate the $\alpha$ but the effect of the former one was better; 3) Pre-screening for rapidly evolving genes performed better than using mammalian-specific genes.

The third category applied the ideas from either (or both) of these two methods to generate a new tool and make new discoveries. Welch et al. (2006) introduced an improvement for the MK estimate α because the conventional way of doing so might be subjected to biases such as reflecting demographic changes rather than adaptive substitution; Using between-locus variation yielded contradictory results; Estimation from the same model organisms have also varied widely [52]. Welch et al. (2006) used three sets of $d_S$ and $d_N$ measurements: the total divergence between *D. yakuba* and *D. simulans*, the total divergence between *D. melanogaster* and *D. simulans*, and the divergence along the *D. simulans* lineage alone, estimated using PAML.

The last category is a pure theoretical comparison based on simulations. Zhai et al. (2009) investigated the statistical power of three neutrality tests for comparative data [124]: the HKA test [125], the MK test [12], and the $d_N/d_S$ likelihood ratio test [14,40], along with two tests based on population genetic data. Using forward simulations, this paper shows that: 1) the most important role of the MK test in population genetics might perhaps be to test for negative selection, whereas other tests should be used to detect positive selection. 2) Under the assumption of a fixed-position model, the $d_N/d_S$ ratio has more power to detect recurrent positive selection than any of the tests which use population genetic data. 3) If more species are included and/or if the divergence time is longer, the power of $d_N/d_S$ ratio tests increases.

***Supplementary note of theoretical background***

Kimura (1957) has shown the fixation probability $f_1$ of an advantage mutant is

$$f_1 = \frac{1-e^{-S_1}}{1-e^{-2NS_1}}$$

where $N$ is population size of a diploid population [35]. The relative fitness of three alleles are $1+S_1$, $1+S_1/2$ and 1 ($S_1>0$).

Hence, the fixation probability $f_2$ of a deleterious mutant could be derived as

$$f_2 = \frac{1-e^{\frac{-S_2}{1+S_2}}}{1-e^{\frac{-2NS_2}{1+S_2}}}$$

The relative fitness of three alleles are $1+S_2$, $1+S_2/2$ and 1 ($S_2 < 0$). Given $|S_2| << 1$, the above formula may be simplified as

$$f_2 \approx \frac{1-e^{-S_2}}{1-e^{-2NS_2}}$$

Hence, $f \approx (1 - e^{-s}) / (1 - e^{-2Ns})$ is the fixation probability of a mutation with a selective coefficient $s$ that can be > 0 (denoted by $s_1$) or < 0 ($s_2$) and half the value in heterozygotes.

On the other hand, the fixation probability $f$ of a neutral mutant is $1/2N$ [33,126]. Then, Ka/Ks = $f$ / $(1/2N)$ = $2Nf$. The Ka/Ks of total sequences could be the weighted sum of the three parts (neutral, advantage and deleterious) as shown in Eq. (1) in the end.

***The results of branch-site model***

The results of using branch-site model of PAML are displayed in Fig. S1 and Table S2-S3. The result in Fig. S1 is similar to Fig. 2 that used site model of PAML. Likewise, similar to Table 1, Table S2 displayed the overlaps between PAML and MK tests with the threshold P < 0.05. In *Drosophila* species, 293 and 186 genes (5.40% and 3.43%) were called by PAML (branch-site model) and MK test, respectively. Only 19 genes are called by both tests, though it is significantly higher than the expected 10.05. Among the 12975 genes in *Arabidopsis*, only a few genes overlap between two tests (21 genes, 0.16% of the genome). Results of the lower stringency (P < 0.224) similar to Table 2 are given in Table S3.

To reduce false negatives, 20-30 neighboring genes were merged into "supergene". In *Drosophila*, 16 out of the 200 supergenes are detected by both tests (Table S2), but their overlap is still almost at random (8.00%, slightly less than the expected 10.08%). In *Arabidopsis*, the two tests also appear to be not correlated (observed 2.20% vs. the expected 3.87%). In addition, similar to the result of site model (see Table 1), the overlap analysis of component genes did not give a satisfactory result.

***Supplementary note of Figure S2***

The 16 scenarios modeled in Fig. S2 represent a sample of the numerous possibilities for selection to vary. The sample is designed to answer the following questions: How often can fluctuating positive selection be detected by PAML and by MK, respectively? The question, partially answered in previous publications, is not the main focus and is relevant here in the context of the second question: How often are the results concordant, or non-concordant when both the MK and PAML tests have sufficient powers? In all scenarios, 3000 genes were monitored and, in 1000 of them, positive selection is operative. Hence, a fully powerful test should yield an estimate of 1000 genes under positive selection. The 16 scenarios fall into two groups, organized into section A and section B.

In Section A, both tests have at least some power to detect selection. The criteria for inclusion in Sec. A is that i) the more powerful test (always PAML in this section) detects > 200 cases out of 1000; and ii) the less powerful test detects > 20 cases. The concordance between the MK and PAML tests can be seen in two different ways.

First, in "overlap" column, the overlap between the two tests is always much larger than the null expectation when the two tests are uncorrelated. Second, in the concordance column, the ratio of the observed overlap to the maximal overlap is defined as the concordance. Note that the number of cases detected in the less powerful test is the maximal concordance. (The expected value for random association is given in the parentheses.)

Section A shows that the observed concordance is generally > 90% when both tests are sufficiently powerful. Even in the two least concordant sets of results (A7 and A8), the observed concordance is still much higher than the expected value based on random association.

Section B shows the scenarios whereby either or both tests have no power in detecting positive selection. The results are of less interest here. In B6 and B7, MK is not expected to detect selection since neither scenario allows selection in the relevant branch. In B8, there is no selection in the entire phylogeny, and neither rest should yield positive results.

In summary, permitting positive selection to fluctuate, we still reach the conclusion that the MK and PAML tests are highly concordant in detecting positive selection. We should further note that if we permit positive selection to fluctuate from time to time, it makes more sense if we should also permit negative selection to fluctuate. There are many biological reasons underlying the fluctuation of negative selection, which is the subject of the companion study [55].

***On fluctuating negative selection affecting the concordance between MK and PAML***

Eq. (2) shows that $q$ can be estimated by Pa/Ps (~ 1- $q$) from the polymorphism data within each species. Thus, a simple test of the constancy of $q$ is to compare the Pa/Ps ratio in each species of interest. For example, between *A. thaliana* and *A. lyrata* (*subsp. lyrata*), the Pa/Ps ratio is 0.142 and 0.248, and the Ka/Ks ratio is 0.215 (Table S5). Clearly, the strength of negative selection has changed in this short time span. In this case, the MK test would reach opposite conclusions depending on whether the polymorphism data used come from *A. thaliana* or *A. lyrata*. Obviously, if two MK tests do not agree, MK should not be expected to agree with the PAML test.

How fluctuating negative selection would affect the PAML tests is more complicated since PAML is a collection of tests, each with a set of assumptions about how positive and negative selection operates [7,18,36,41]. Let us consider the site model in Fig. 1 of the main text. If the red branch in Fig. 1a and 1b experience less negative selection than the remaining branches of the phylogeny, then the Pa/Ps ratio would be relatively high, leading to the call of positive selection on the branch. In this scenario, MK would not call positive selection since the polymorphism would have the same high Pa/Ps ratio.

In short, most scenarios of fluctuating negative selection would bias the calls of positive selection by either test, but in different manners.

***Factors driving the fluctuation in negative selection***

In light of the current report, we now outline a path forward that is elaborated in the companion study [55]. In taking the new path, we are concerned with measuring the fluctuation in negative selection without addressing the many factors underlying the fluctuation. These factors may include environment, genetic background, nucleotide substitution pattern, population size, population subdivision, and so on. The concept of the effective population size, $N_e$, incorporates many (but not all) of such factors. For example, if the actual population size fluctuates, $N_e$ can often be expressed as the harmonic mean of $N$ through time. Nevertheless, $N_e$ could not possibly be the only factor for the variation in the strength of negative selection. A simple example is as follows: Let $s$ be - 0.005. In that case, $N_e$ = 1000, 10000, or higher does not matter because the fixation probability is almost zero if $2N_e s < -10$. By this reasoning, $N_e$ fluctuation may not affect the long term molecular evolutionary rate. In short, while the variation in Pa/Ps can be documented, the underlying causes of such variation may be fairly complex, and those causes are not germane to the current study.

**Supplementary References**

60. dos Santos G, Schroeder AJ, Goodman JL *et al.* FlyBase: Introduction of the *Drosophila melanogaster* Release 6 reference genome assembly and large-scale migration of genome annotations. *Nucleic Acids Res* 2014;**43**:D690–7.

61. Waterhouse RM, Tegenfeldt F, Li J *et al.* OrthoDB: A hierarchical catalog of animal, fungal and bacterial orthologs. *Nucleic Acids Res* 2013;**41**:D358–65.

62. Edgar RC. MUSCLE: Multiple sequence alignment with high accuracy and high throughput. *Nucleic Acids Res* 2004;**32**:1792–7.

63. Suyama M, Torrents D, Bork P. PAL2NAL: Robust conversion of protein sequence alignments into the corresponding codon alignments. *Nucleic Acids Res* 2006;**34**:W609–12.

64. Hoskins RA, Carlson JW, Wan KH *et al.* The Release 6 reference sequence of the *Drosophila melanogaster* genome. *Genome Res* 2015;**25**:445–58.

65. Hu TT, Eisen MB, Thornton KR *et al.* A second-generation assembly of the *Drosophila simulans* genome provides new insights into patterns of lineage-specific divergence. *Genome Res* 2013;**23**:89–98.

66. Pool JE, Corbett-Detig RB, Sugino RP *et al.* Population genomics of Sub-Saharan *Drosophila melanogaster*: African diversity and non-African admixture. *PLoS Genet* 2012;**8**:e1003080.

67. Signor SA, New FN, Nuzhdin S. A large panel of *Drosophila simulans* reveals an abundance of common variants. *Genome Biol Evol* 2018;**10**:189–206.

68. Rogers RL, Cridland JM, Shao L *et al.* Landscape of standing variation for tandem duplications in *Drosophila yakuba* and *Drosophila simulans*. *Mol Biol Evol* 2014;**31**:1750–66.

69. Schrider DR, Ayroles J, Matute DR *et al.* Supervised machine learning reveals introgressed loci in the genomes of *Drosophila simulans* and *D. sechellia*. *PLoS Genet* 2018;**14**:e1007341.

70. Li H, Durbin R. Fast and accurate short read alignment with Burrows-Wheeler transform. *Bioinformatics* 2009;**25**:1754–60.

71. Li H, Handsaker B, Wysoker A *et al.* The sequence alignment/map format and SAMtools. *Bioinformatics* 2009;**25**:2078–9.

72. DePristo MA, Banks E, Poplin R *et al.* A framework for variation discovery and genotyping using next-generation DNA sequencing data. *Nat Genet* 2011;**43**:491–8.

73. Narasimhan V, Danecek P, Scally A *et al.* BCFtools/RoH: A hidden Markov model approach for detecting autozygosity from next-generation sequencing data. *Bioinformatics* 2016;**32**:1749–51.

74. Danecek P, Auton A, Abecasis G *et al.* The variant call format and VCFtools. *Bioinformatics* 2011;**27**:2156–8.

75. Goodstein DM, Shu S, Howson R *et al.* Phytozome: A comparative platform for green plant genomics. *Nucleic Acids Res* 2012;**40**:1178–86.

76. Lamesch P, Berardini TZ, Li D *et al.* The *Arabidopsis* Information Resource (TAIR): Improved gene annotation and new tools. *Nucleic Acids Res* 2012;**40**:1202–10.

77. Hu TT, Pattyn P, Bakker EG *et al.* The *Arabidopsis lyrata* genome sequence and the basis of rapid genome size change. *Nat Genet* 2011;**43**:476–83.

78. Slotte T, Hazzouri KM, Ågren JA *et al.* The *Capsella rubella* genome and the genomic consequences of rapid mating system evolution. *Nat Genet* 2013;**45**:831–5.

79. Lee C-R, Wang B, Mojica JP *et al.* Young inversion with multiple linked QTLs under selection in a hybrid zone. *Nat Ecol Evol* 2017;**1**:119.

80. Remm M, Storm CEV, Sonnhammer ELL. Automatic clustering of orthologs and in-paralogs from pairwise species comparisons. *J Mol Biol* 2001;**314**:1041–52.

81. Mattila TM, Tyrmi J, Pyhäjärvi T *et al.* Genome-wide analysis of colonization history and concomitant selection in *Arabidopsis lyrata*. *Mol Biol Evol* 2017;**34**:2665–77.

82. Josephs EB, Lee YW, Stinchcombe JR *et al.* Association mapping reveals the role of purifying selection in the maintenance of genomic variation in gene expression. *Proc Natl Acad Sci U S A* 2015;**112**:15390–5.

83. Ågren JA, Wang W, Koenig D *et al.* Mating system shifts and transposable element evolution in the plant genus Capsella. *BMC Genomics* 2014;**15**:602.

84. Sayers EW, Beck J, Brister JR *et al.* Database resources of the National Center for Biotechnology Information. *Nucleic Acids Res* 2019;**48**:D9–16.

85. Schneider VA, Graves-Lindsay T, Howe K *et al.* Evaluation of GRCh38 and de novo haploid genome assemblies demonstrates the enduring quality of the reference assembly. *Genome Res* 2017;**27**:849–64.

86. Zhou X, Wang B, Pan Q *et al.* Whole-genome sequencing of the snub-nosed monkey provides insights into folivory and evolutionary history. *Nat Genet* 2014;**46**:1303–10.

87. Palesch D, Bosinger SE, Tharp GK *et al.* Sooty mangabey genome sequence provides insight into AIDS resistance in a natural SIV host. *Nature* 2018;**553**:77–81.

88. Locke DP, Hillier LW, Warren WC *et al.* Comparative and demographic analysis of orang-utan genomes. *Nature* 2011;**469**:529–33.

89. Hughes JF, Skaletsky H, Pyntikova T *et al.* Conservation of Y-linked genes during human evolution revealed by comparative sequencing in chimpanzee. *Nature* 2005;**437**:100–3.

90. Prüfer K, Munch K, Hellmann I *et al.* The bonobo genome compared with the chimpanzee and human genomes. *Nature* 2012;**486**:527–31.

91. Scally A, Dutheil JY, Hillier LW *et al.* Insights into hominid evolution from the gorilla genome sequence. *Nature* 2012;**483**:169–75.

92. Carbone L, Alan Harris R, Gnerre S *et al.* Gibbon genome and the fast karyotype evolution of small apes. *Nature* 2014;**513**:195–201.

93. Warren WC, Jasinska AJ, García-Pérez R *et al.* The genome of the vervet (*Chlorocebus aethiops sabaeus*). *Genome Res* 2015;**25**:1921–33.

94. Zimin A V, Cornish AS, Maudhoo MD *et al.* A new rhesus macaque assembly and annotation for next-generation sequencing analyses. *Biol Direct* 2014;**9**:20.

95. Rogers J, Raveendran M, Harris RA *et al.* The comparative genomics and complex population history of *Papio baboons*. *Sci Adv* 2019;**5**:eaau6947.

96. Yu L, Wang GD, Ruan J *et al.* Genomic analysis of snub-nosed monkeys (*Rhinopithecus*) identifies genes and processes related to high-altitude adaptation. *Nat Genet* 2016;**48**:947–52.

97. Worley KC, Warren WC, Rogers J *et al.* The common marmoset genome provides insight into primate biology and evolution. *Nat Genet* 2014;**46**:850–7.

98. Emms DM, Kelly S. OrthoFinder: Phylogenetic orthology inference for comparative genomics. *Genome Biol* 2019;**20**:238.

99. Buchfink B, Xie C, Huson DH. Fast and sensitive protein alignment using DIAMOND. *Nat Methods* 2015;**12**:59–60.

100. Suzuki Y, Gojobori T, Kumar S. Methods for incorporating the hypermutability of CpG dinucleotides in detecting natural selection operating at the amino acid sequence level. *Mol Biol Evol* 2009;**26**:2275–84.

101. Walser JC, Furano A V. The mutational spectrum of non-CpG DNA varies with CpG content. *Genome Res* 2010;**20**:875–82.

102. Gibbs RA, Boerwinkle E, Doddapaneni H *et al.* A global reference for human genetic variation. *Nature* 2015;**526**:68–74.

103. Sudmant PH, Rausch T, Gardner EJ *et al.* An integrated map of structural variation in 2,504 human genomes. *Nature* 2015;**526**:75–81.

104. Prado-Martinez J, Sudmant PH, Kidd JM *et al.* Great ape genetic diversity and population history. *Nature* 2013;**499**:471–5.

105. Liu Z, Tan X, Orozco-terWengel P *et al.* Population genomics of wild Chinese rhesus macaques reveals a dynamic demographic history and local adaptation, with implications for biomedical research. *Gigascience* 2018;**7**:giy106.

106. Zhang G, Li B, Li C *et al.* Comparative genomic data of the Avian Phylogenomics Project. *Gigascience* 2014;**3**:26.

107. Zhang G, Li C, Li Q *et al.* Comparative genomics reveals insights into avian genome evolution and adaptation. *Science* 2014;**346**:1311–20.

108. Jarvis ED, Mirarab S, Aberer AJ *et al.* Whole-genome analyses resolve early branches in the tree of life of modern birds. *Science* 2014;**346**:1320–31.

109. Li S, Li B, Cheng C *et al.* Genomic signatures of near-extinction and rebirth of the crested ibis and other endangered bird species. *Genome Biol* 2014;**15**:557.

110. Gibb GC, Kennedy M, Penny D. Beyond phylogeny: Pelecaniform and ciconiiform birds, and long-term niche stability. *Mol Phylogenet Evol* 2013;**68**:229–38.

111. Slack KE, Jones CM, Ando T *et al.* Early penguin fossils, plus mitochondrial genomes, calibrate avian evolution. *Mol Biol Evol* 2006;**23**:1144–55.

112. Haddrath O, Baker AJ. Complete mitochondrial DNA genome sequences of extinct birds: Ratite phylogenetics and the vicariance biogeography hypothesis. *Proc R Soc B Biol Sci* 2001;**268**:939–45.

113. Haller BC, Messer PW. SLiM 3: Forward genetic simulations beyond the Wright-Fisher model. *Mol Biol Evol* 2019;**36**:632–7.

114. Kimura M. A simple method for estimating evolutionary rates of base substitutions through comparative studies of nucleotide sequences. *J Mol Evol* 1980;**16**:111–20.

115. Haller BC, Messer PW. SLiM: An evolutionary simulation framework. 2016.

116. Jukes TH, Cantor CR. Evolution of protein molecules. *Mamm protein Metab* 1969;**3**:132.

117. Zhao H, Sun Z, Wang J *et al.* CrossMap: a versatile tool for coordinate conversion between genome assemblies. *Bioinformatics* 2014;**30**:1006–7.

118. Cingolani P, Platts A, Wang LL *et al.* A program for annotating and predicting the effects of single nucleotide polymorphisms, SnpEff. *Fly* 2012;**6**:80–92.

119. Zeng K, Shi S, Fu Y-X *et al.* Statistical tests for detecting positive selection by utilizing high frequency variants. *Genetics* 2006;**174**:1–33.

120. Zeng K, Mano S, Shi S *et al.* Comparisons of site- and haplotype-frequency methods for detecting positive selection. *Mol Biol Evol* 2007;**24**:1562–74.

121. Zeng K, Shi S, Wu CI. Compound tests for the detection of hitchhiking under positive selection. *Mol Biol Evol* 2007;**24**:1898–908.

122. Feldmeyer B, Greshake B, Funke E *et al.* Positive selection in development and growth rate regulation genes involved in species divergence of the genus Radix. *BMC Evol Biol* 2015;**15**:164.

123. Wong A, Turchin MC, Wolfner MF *et al.* Evidence for positive selection on *Drosophila melanogaster* seminal fluid protease homologs. *Mol Biol Evol* 2007;**25**:497–506.

124. Zhai W, Nielsen R, Slatkin M. An investigation of the statistical power of neutrality tests based on comparative and population genetic data. *Mol Biol Evol* 2009;**26**:273–83.

125. Hudson RR, Kreitman M, Aguadé M. A test of neutral molecular evolution based on nucleotide data. *Genetics* 1987;**116**:153–9.

126. Cormack RM, Hartl DL, Clark AG. *Principles of Population Genetics*. Sunderland (MA): Sinauer Associates, 1990.

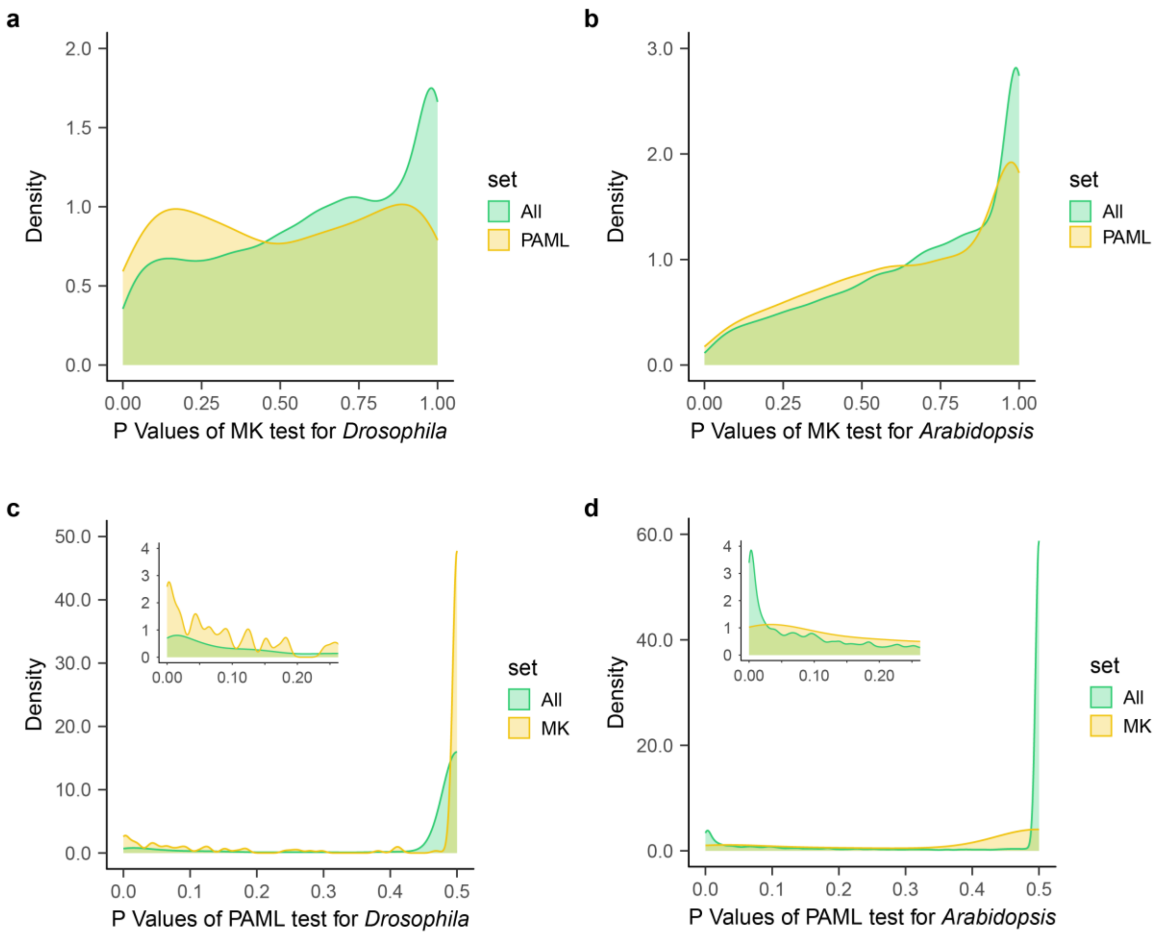


**Figure S1. P value distributions of MK and PAML tests (the branch-site model).** In panels **a** and **b**, the P values of the MK test for all genes are shown in green, and that for genes selected by PAML test are shown in yellow. In panels **c** and **d**, the green distributions represent the P values of the PAML test for all genes, and the yellow distributions the genes selected by MK test.

**a**

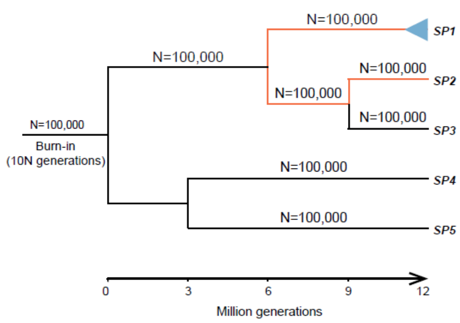


**b**

| Group | Branch | Ns=0 | Ns=-1 | Ns=-10 | Ns=-100 | Ns=5 |
|---|---|---|---|---|---|---|
| 1 | blue | 5% | 5% | 10% | 80% | 0% |
| | black | 5% | 5% | 10% | 75% | 5% |
| 2 | blue | 10% | 5% | 5% | 80% | 0% |
| | black | 10% | 5% | 5% | 75% | 5% |
| 3 | blue | 5% | 5% | 5% | 85% | 0% |
| | black | 5% | 5% | 5% | 75% | 10% |

**c**

| Section A model | PAML | Observed overlap (Expected) | MK | Observed concordance (Expected) |
|---|---|---|---|---|
| A1 | 1006 | 35 (12.4) | 37 | 94.6% (33.5%) |
| | 1026 | 56 (22.6) | 66 | 84.8% (34.2%) |
| | 1018 | 121 (41.1) | 121 | 100.0% (33.9%) |
| A2 | 986 | 40 (13.8) | 42 | 95.2% (32.9%) |
| | 982 | 57 (21.9) | 67 | 85.1% (32.7%) |
| | 1016 | 115 (38.9) | 115 | 100.0% (33.9%) |
| A3 | 1006 | 38 (13.4) | 40 | 95.0% (33.5%) |
| | 1023 | 48 (19.8) | 58 | 82.8% (34.1%) |
| | 1017 | 102 (34.6) | 102 | 100.0% (33.9%) |
| A4 | 1006 | 35 (12.4) | 37 | 94.6% (33.5%) |
| | 1026 | 70 (27.4) | 80 | 87.5% (34.2%) |
| | 1018 | 97 (32.9) | 97 | 100.0% (33.9%) |
| A5 | 1006 | 37 (13.1) | 39 | 94.9% (33.5%) |
| | 1022 | 55 (22.1) | 65 | 84.6% (34.1%) |
| | 1018 | 185 (62.8) | 185 | 100.0% (33.9%) |
| A6 | 990 | 43 (14.9) | 45 | 95.6% (33.0%) |
| | 994 | 62 (24.5) | 74 | 83.8% (33.1%) |
| | 1016 | 114 (38.6) | 114 | 100.0% (33.9%) |
| A7 | 765 | 16 (5.9) | 23 | 69.6% (25.5%) |
| | 720 | 22 (9.8) | 41 | 53.7% (24.0%) |
| | 884 | 50 (16.8) | 57 | 87.7% (29.5%) |
| A8 | 217 | 13 (3.4) | 47 | 27.7% (7.2%) |
| | 233 | 12 (5.7) | 73 | 16.4% (7.8%) |
| | 259 | 26 (9.9) | 115 | 22.6% (8.6%) |

| Section B model | PAML | Observed overlap (Expected) | MK |
|---|---|---|---|
| B1 | 1001 | 3 (1.67) | 5 |
| | 1010 | 6 (5.39) | 16 |
| | 1012 | 18 (6.07) | 18 |
| B2 | 1004 | 7 (3.01) | 9 |
| | 1025 | 9 (6.49) | 19 |
| | 1017 | 16 (5.42) | 16 |
| B3 | 8 | 0 (0.03) | 11 |
| | 32 | 0 (0.21) | 20 |
| | 20 | 1 (0.09) | 13 |
| B4 | 10 | 0 (0.10) | 29 |
| | 27 | 0 (0.34) | 38 |
| | 20 | 0 (0.42) | 63 |
| B5 | 22 | 1 (0.29) | 40 |
| | 48 | 2 (0.98) | 61 |
| | 24 | 0 (0.71) | 89 |
| B6 | 181 | 0 (/) | 4 |
| | 180 | 2 (/) | 15 |
| | 234 | 0 (/) | 0 |
| B7 | 210 | 0 (/) | 2 |
| | 202 | 0 (/) | 12 |
| | 333 | 0 (/) | 3 |
| B8 | 9 | 0 (//) | 3 |
| | 39 | 0 (//) | 15 |
| | 27 | 0 (//) | 0 |

**Figure S2. Diagram of simulations for PAML and MK tests. a**, the tree shows the simulation process. MK test uses polymorphisms (indicated by the blue triangles) to detect positive selection along the red branches. PAML test is used to detect positive selection along whole lineages including red branches. **b** and **c**, 16 simulation models that positive selection operates on chosen branches of the phylogenetic tree of panel **a** and results of PAML and MK tests. The parameter values are given in **b** and the results are given in **c**.

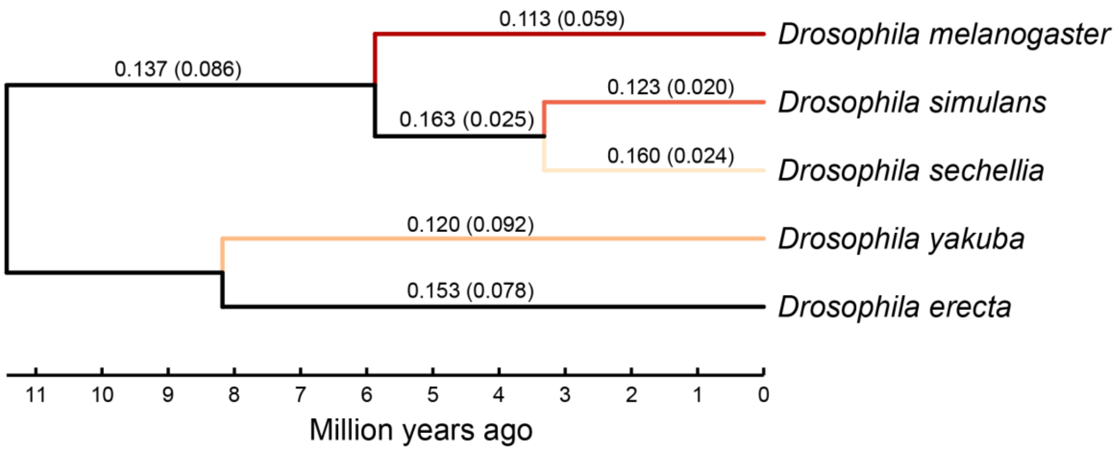


**Figure S3. Ka/Ks ratios (and Ks) along each branch of the *Drosophila melanogaster* subgroup phylogeny.** The Ka/Ks ratios of colored branches are the lineage-specific ratios shown in Fig. 3a.

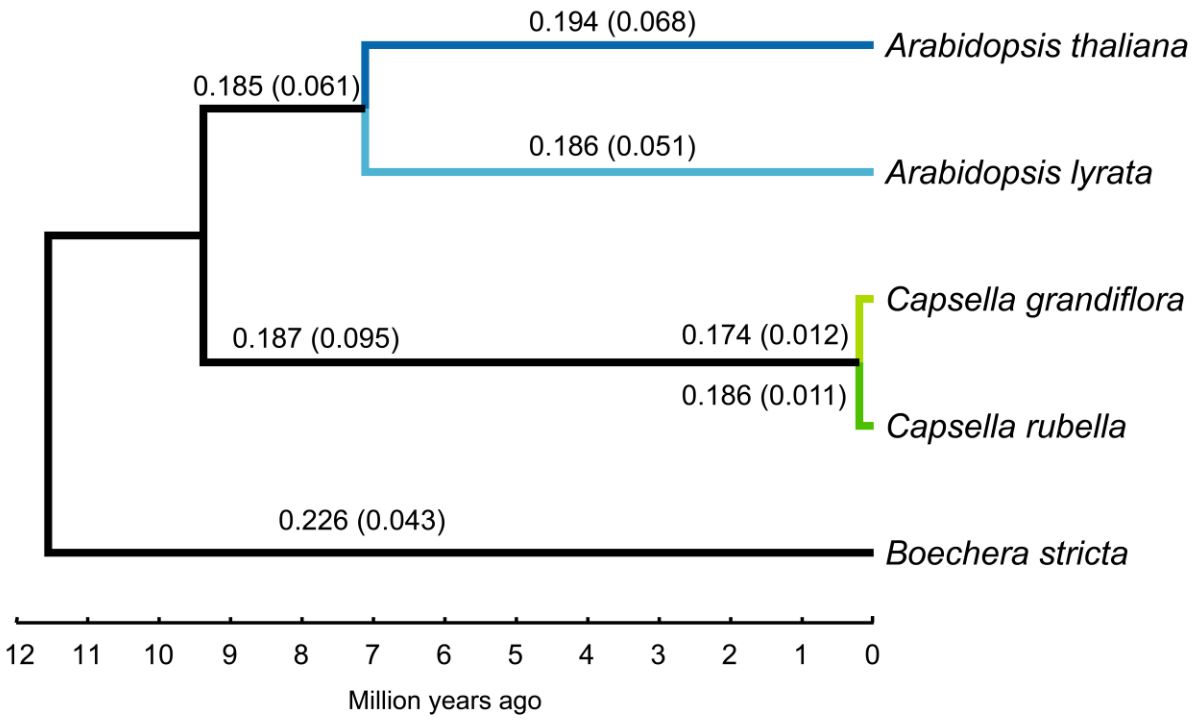


**Figure S4. Ka/Ks ratios (and Ks) along each branch of the *Arabidopsis thaliana* group phylogeny.** The Ka/Ks ratios of colored branches are the lineage-specific ratios shown in Fig. 3b.

**Table S1. Proportion of adaptively evolving genes identified by two tests ($P < 0.05$)**

**(Same as Table 1, but genes were merged into supergenes by ontology)**

| Gene Category | MK | PAML (site model) | Expected overlap | Observed overlap |
|---|---|---|---|---|
| ***Drosophila*** | | | | |
| Supergenes[a] | **31.52%** (58/184) | **14.67%** (27/184) | **4.62%** | **8.70%** (16/184) |
| Component genes[b] | **6.01%** (51/849) | **7.54%** (36/477) | **0.45%** | **0.00%** (0/306) |
| ***Arabidopsis*** | | | | |
| Supergenes[a] | **10.57%** (48/454) | **19.38%** (88/454) | **2.05%** | **2.42%** (11/454) |
| Component genes[b] | **4.46%** (45/1008) | **7.19%** (184/2556) | **0.32%** | **0.29%** (1/341) |

[a] Supergenes are the concatenations of genes of the same ontology.

[b] Component genes are individual genes within supergenes that have passed the MK and/or PAML tests.

**Table S2. Proportion of adaptively evolving genes identified by two tests (P < 0.05)**

**(Same as Table 1 but using the PAML branch-site model)**

| Gene Category | MK | PAML (branch-site) | Expected overlap | Observed overlap |
|---|---|---|---|---|
| ***Drosophila*** | | | | |
| Individual genes | **3.43%** (186/5425) | **5.40%** (293/5425) | **0.19%** (10.05/5425) | **0.35%** (19/5425) |
| Supergenes[a] | **56.00%** (112/200) | **18.00%** (36/200) | **10.08%** | **8.00%** (16/200) |
| Component genes[b] | **5.04%** (158/3132) | **9.41%** (92/978) | **0.47%** | **1.76%** (8/455) |
| ***Arabidopsis*** | | | | |
| Individual genes | **1.12%** (145/12975) | **10.02%** (1300/12975) | **0.12%** (14.53/12975) | **0.16%** (21/12975) |
| Supergenes | **8.20%** (41/500) | **47.20%** (236/500) | **3.87%** | **2.20%** (11/500) |
| Component genes | **3.62%** (38/1048) | **12.24%** (750/6129) | **0.44%** | **1.36%** (4/295) |

[a] Supergenes are concatenations of 20-30 neighboring genes by physical location.

[b] Component genes are individual genes within supergenes that have passed the MK and/or PAML tests.

**Table S3. Proportion of adaptively evolving sites identified by two tests ($P^2 < 0.05$, i.e. $P < 0.224$)**

**(Same as Table 2 but using the PAML branch-site model)**

| | **MK** | **MK-PAML overlap** | **PAML (branch-site)** | **Total** |
|---|---|---|---|---|
| ***Drosophila*** | | | | |
| No. of genes | 824 | 127[d] | 530 | 5425 |
| Expected overlap | / | 80.50 | / | / |
| Proportion of adaptive changes by MK[a] | 0.69 | 0.65 | 0.31 | 0.26 |
| No. of adaptive sites per gene by MK (A1) | 14.98 | 20.22 | 6.24 | 2.84 |
| No. of adaptive sites per gene by PAML (A2)[b] | 8.33 | 23.96 | 22.40 | 5.02 |
| No. of adaptive sites per gene by PAML (A2')[c] | 1.95 | 9.69 | 9.53 | 1.23 |
| ***Arabidopsis*** | | | | |
| No. of genes | 1014 | 233[e] | 1172 | 12975 |
| Expected overlap | / | 193.89 | / | / |
| Proportion of adaptive changes by MK | 0.69 | 0.69 | 0.06 | 0.04 |
| No. of adaptive sites per gene by MK (A1) | 19.36 | 24.07 | 1.50 | 0.84 |
| No. of adaptive sites per gene by PAML (A2) | 4.48 | 9.94 | 7.78 | 3.33 |
| No. of adaptive sites per gene by PAML (A2') | 2.72 | 9.00 | 7.69 | 2.06 |

[a] Proportion of adaptive changes is done using Shapiro et al. (2007)'s method of correction [3].

[b] A2 is based on PAML-M2a model.

[c] A2' is based on PAML-BEB model.

[d] $P < 10^{-7}$ by Fisher's exact test, given 80.5 as the expected value.

[e] $P < 10^{-10}$ by Fisher's exact test, given 193.9 as the expected value.

**Table S4. Summary of non-synonymous and synonymous mutations in *Arabidopsis***

| | *A. thaliana* | *A. lyrata petraea* | *A. lyrata lyrata* | *C. grandiflora* | *C. rubella* |
|---|---|---|---|---|---|
| **Sample Size** | 1,135 | 22 | 6 | 8 | 12 |
| **Expect non-synonymous sites** | 12,112,293 | 12,111,370 | 12,111,370 | 12,110,726 | 12,110,695 |
| **Expect synonymous sites** | 4,319,514 | 4,320,443 | 4,320,443 | 4,321,093 | 4,321,124 |
| **Polymorphism (all sites)** | | | | | |
| Non-synonymous | 235,079 | 167,802 | 26,282 | 128,800 | 24,021 |
| Synonymous | 241,506 | 259,432 | 34,448 | 267,800 | 43,743 |
| Pa/Ps ratio | 0.347 | 0.231 | 0.272 | 0.172 | 0.196 |
| **Polymorphism (Freq. <= 0.2)** | | | | | |
| Non-synonymous | 211,817 | 112,563 | 8,344 | 87,378 | 15,084 |
| Synonymous | 186,781 | 141,987 | 8,619 | 150,114 | 24,570 |
| Pa/Ps ratio | 0.404 | 0.283 | 0.345 | 0.208 | 0.219 |
| **Polymorphism (Freq. > 0.2)** | | | | | |
| Non-synonymous | 23,262 | 55,239 | 17,938 | 41,422 | 8,937 |
| Synonymous | 54,725 | 117,445 | 25,829 | 117,686 | 19,173 |
| Pa/Ps ratio | 0.152 | 0.168 | 0.248 | 0.126 | 0.166 |
| **Lineage-specific Divergence** | | | | | |
| Non-synonymous | 159,662 | 114,613 | 114,613 | 24,538 | 25,841 |
| Synonymous | 282,693 | 213,754 | 213,754 | 49,892 | 49,296 |
| Ka/Ks ratio | 0.194 | 0.186 | 0.186 | 0.174 | 0.186 |

**Table S5. Summary of non-synonymous and synonymous mutations in *Drosophila***

| | *D. melanogaster* | *D. simulans* | *D. sechellia* | *D. yakuba* |
|---|---|---|---|---|
| **Sample Size** | 99 | 170 | 41 | 20 |
| **Expect non-synonymous sites** | 4,881,646 | 4,881,400 | 4,881,820 | 4,880,463 |
| **Expect synonymous sites** | 1,783,805 | 1,784,051 | 1,783,631 | 1,784,988 |
| **Polymorphism (all sites)** | | | | |
| Non-synonymous | 67,144 | 27,662 | 3,859 | 42,160 |
| Synonymous | 173,366 | 136,734 | 6,313 | 174,420 |
| Pa/Ps ratio | 0.142 | 0.074 | 0.223 | 0.242 |
| **Polymorphism (Freq. <= 0.2)** | | | | |
| Non-synonymous | 59,855 | 17,810 | 1,858 | 33,564 |
| Synonymous | 126,189 | 78,502 | 2,800 | 116,240 |
| Pa/Ps ratio | 0.173 | 0.083 | 0.242 | 0.106 |
| **Polymorphism (Freq. > 0.2)** | | | | |
| Non-synonymous | 7,289 | 9,852 | 2,001 | 8,596 |
| Synonymous | 47,177 | 58,232 | 3,513 | 58,180 |
| Pa/Ps ratio | 0.056 | 0.062 | 0.208 | 0.054 |
| **Lineage-specific Divergence** | | | | |
| Non-synonymous | 32,274 | 12,191 | 18,862 | 53,684 |
| Synonymous | 101,019 | 35,655 | 42,419 | 154,885 |
| Ka/Ks ratio | 0.113 | 0.123 | 0.160 | 0.120 |

**Table S6. Summary of non-synonymous and synonymous mutations in primates**

| Species (subspecies or population) | *n*[a] | Pa/Ps ratio: Average heterozygosity[b] | Pa/Ps ratio: Polymorphism (all sites) | Pa/Ps ratio: Polymorphism (no singletons) | Ka/Ks[c] |
|---|---|---|---|---|---|
| ***Homo*** | | | | | |
| *H. sapiens* (Han Chinese) | 106 | 0.390±0.022 | 0.520 | 0.400 | 0.329±0.013 |
| *H. sapiens* (Yoruba population) | 107 | 0.378±0.015 | 0.449 | 0.382 | |
| ***Pan*** | | | | | |
| *P. paniscus* | 13 | 0.437±0.024 | 0.419 | 0.384 | 0.321±0.013 |
| *P. troglodytes verus* | 4 | 0.429±0.017 | 0.397 | 0.353 | 0.327±0.013 |
| *P. troglodytes ellioti* | 10 | 0.393±0.014 | 0.373 | 0.344 | |
| *P. troglodytes schweinfurthii* | 6 | 0.368±0.016 | 0.344 | 0.309 | |
| *P. troglodytes troglodytes* | 4 | 0.370±0.011 | 0.338 | 0.295 | |
| ***Gorilla*** | | | | | |
| *G. beringei graueri* | 3 | 0.397±0.022 | 0.356 | 0.341 | - |
| *G. gorilla gorilla* | 23 | 0.360±0.011 | 0.372 | 0.331 | 0.325±0.013 |
| ***Pongo*** | | | | | |
| *P. pygmaeus* | 5 | 0.360±0.009 | 0.320 | 0.298 | - |
| *P. abellii* | 5 | 0.328±0.010 | 0.291 | 0.282 | 0.302±0.013 |
| ***Rhinopithecus*** | | | | | |
| *R. roxellana* | 18 | 0.380±0.028 | 0.355 | 0.332 | 0.301±0.017 |
| *R. bieti* | 20 | 0.393±0.009 | 0.359 | 0.349 | 0.338±0.016 |
| ***Macaca*** | | | | | |
| *M. mulatta lasiotus* | 32 | 0.276±0.009 | 0.275 | 0.245 | 0.310±0.018 |
| *M. mulatta littoralis* | 29 | 0.272±0.007 | 0.270 | 0.244 | |
| *M. mulatta tcheliensis* | 5 | 0.275±0.011 | 0.244 | 0.240 | |
| *M. mulatta brevicaudatus* | 5 | 0.275±0.014 | 0.248 | 0.237 | |
| *M. mulatta mulatta* | 10 | 0.278±0.005 | 0.253 | 0.243 | |

[a] *n*, sample size of each species (subspecies or population).

[b] Average heterozygosity Pa/Ps ratios with standard deviation in *n* individuals of each species or subspecies.

[c] Average Ka/Ks ratios with standard deviation compared to each species in other families. (Between Hominoids and old-world monkeys).
